# Structural and biochemical requirements for secretory component interactions with dimeric Immunoglobulin A

**DOI:** 10.1101/2023.11.09.566401

**Authors:** Sonya Kumar Bharathkar, Beth M. Stadtmueller

## Abstract

Secretory (S) Immunoglobulin (Ig) A is the predominant mucosal antibody that protects host epithelial barriers and promotes microbial homeostasis. SIgA production occurs when plasma cells assemble two copies of monomeric IgA and one joining-chain (JC) to form dimeric (d) IgA, which is bound by the polymeric Ig receptor (pIgR) on the basolateral surface of epithelial cells and transcytosed to the apical surface. There, pIgR is proteolytically cleaved, releasing SIgA, a complex of the dIgA and the pIgR ectodomain, called secretory component (SC). The pIgR’s five Ig-like domains (D1-D5) undergo a conformational change upon binding dIgA, ultimately contacting four IgA heavy chains and the JC in SIgA. Here we report structure-based mutational analysis combined with surface plasmon resonance binding assays that identify key residues in mouse SC D1 and D3 that mediate SC binding to dIgA. Residues in D1 CDR3 are likely to initiate binding whereas residues that stabilize the D1-D3 interface are likely to promote the conformation change and stabilize the final SIgA structure. Additionally, we find that the JC’s three C-terminal residues play a limited role in dIgA assembly but a significant role in pIgR/SC binding to dIgA. Together results inform new models for the intricate mechanisms underlying IgA transport across epithelia and functions in the mucosa.

**KEY POINTS:**

1. The secretory component D1 CDR3 loop plays an important role in binding to dIgA.
2. The formation of SC D1-D3 interface stabilizes SIgA.
3. JC C-terminal residues influence dIgA assembly and mediate pIgR binding to dIgA.

## INTRODUCTION

Secretory Immunoglobin A (SIgA) is a crucial antibody in the mucosal immune system that helps maintain the balance of microorganisms and protect the mucosal barrier (1),(2). The assembly of SIgA begins in the lamina propria, where plasma cells link one Joining chain (JC) and two to five copies of monomeric (m) IgA to form polymeric (p) IgA; in humans the predominant form is dimeric (d) IgA(3),(4),(5). Subsequently, pIgA binds to the polymeric Ig Receptor (pIgR) on the basolateral surface of mucosal epithelial cells and is transcytosed to the apical side(6),(7). Subsequently, proteases cleave the pIgR ectodomain from the cell surface, releasing a complex containing pIgR’s five Ig-like domains (D1-D5), collectively referred to as Secretory Component (SC), bound to the pIgA(8),(9). The SC-pIgA complex, which typically includes dIgA, is referred to as SIgA(10).

Each mIgA consists of two heavy chains (HC) and two light chains (LC) that together form two antigen binding fragments (Fabs) and one constant fragment (Fcα), and thus SIgA contains at least four Fabs and two Fcs (Fig.1). The cryo-EM structures of SIgA(11),(12),(13) and dIgA(11) revealed a remarkably asymmetric molecule with individual mIgAs bent and tilted with respect to each other. This arrangement results in each HC (HC-A-D) adopting a unique structure and unique contacts with other chains, despite sharing the same sequence. The greatest structural variability is found among copies of the HC’s 18-22 residue C-terminal extension, called the tailpiece (Tp) (11),(12),(13). The four Tps (TpA-D) co-fold with JC to form a β-sandwich, in which the Tps closest to JC, TpA and TpC, are disulfide bonded to the JC with their penultimate Cys residue(11),(12),(13),(14). The other two Tps, TpB and TpD, are partially disordered in dIgA, but are ordered in SIgA, indicating a possible role in pIgR binding and successful SIgA formation(11). In addition to contacting the Tps, the JC also contacts HC residues in the Fcα regions, through several β-hairpins, termed *wings*. JC_wing-1_ and JC_wing-2_ contact HC-C and HC-A respectively, and appear to contribute to the bent, asymmetric conformation observed for both dIgA and SIgA. Additionally JC also contacts parts of SC with its C-terminal residues interfacing with SC-D1, and JC_wing-1_ interfacing with SC-D4-D5 (11),(12),(13).

**Figure-1:**
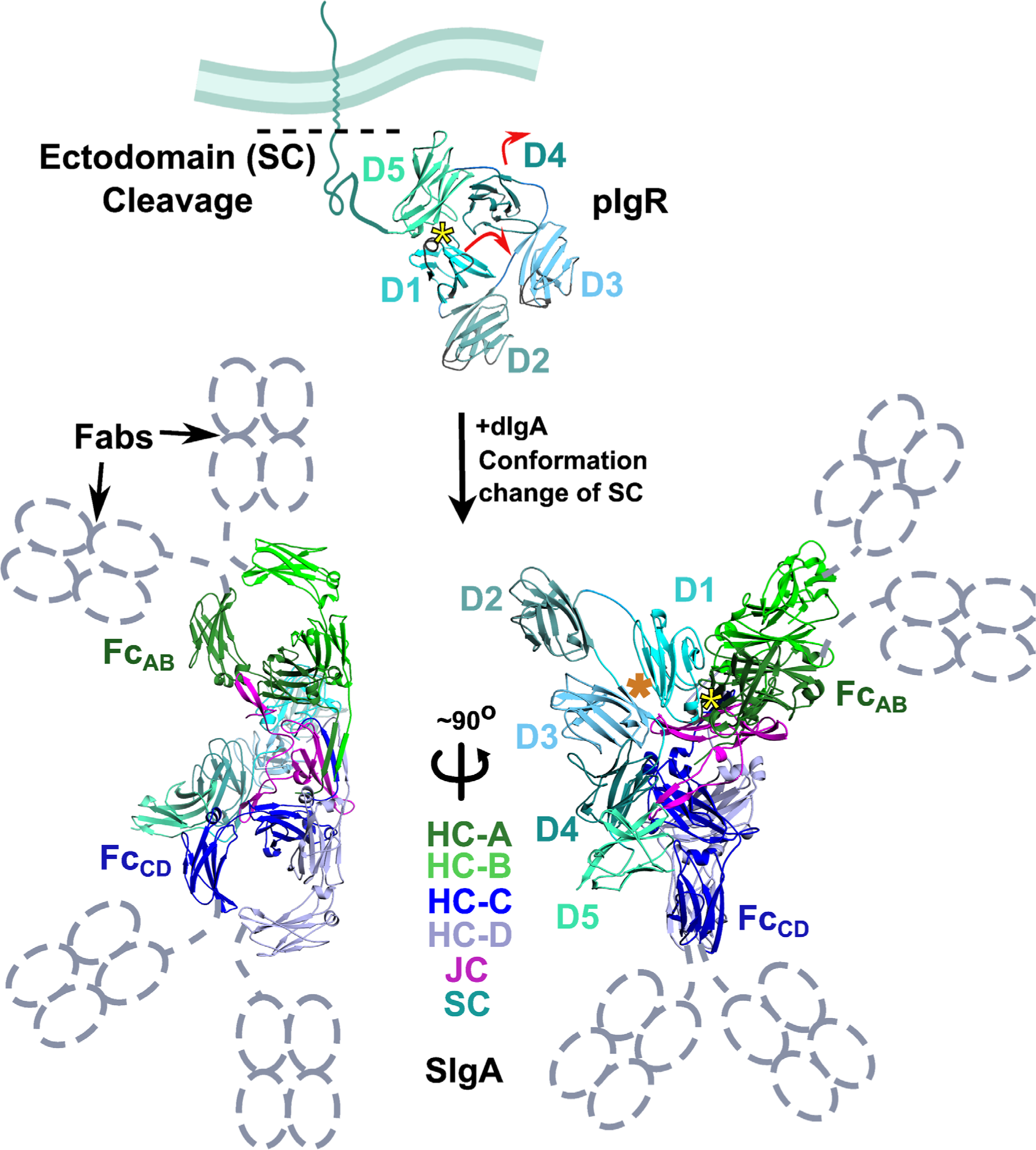
The structure of liganded and unliganded SC. (Top) Cartoon representation of a mouse pIgR ectodomain homology model based on the human SC crystal structure (PDB:5D4K). The pIgR is depicted bound to an epithelial cell membrane with its five constitutive immunoglobin-like domains (D1-D5) adopting a closed, unliganded conformation. Upon encountering dIgA, D1-D5 undergoes conformational change (indicated by red arrows) to form SIgA (PDB:7JG2), which is shown as a cartoon representation in two orientations (Fabs are disordered and represented with dotted schematics in grey). The dimeric SIgA contains four heavy chains (HC-A to D), J-chain (JC), and one Secretory Component (SC). All chains are colored according to the key. The Fc formed by HC-A and HC-B is indicated as Fc_AB_ (green) and the Fc formed by HC-C and HC-D is indicated as Fc_CD_ (of blue). The JC occupies the center of the SIgA complex with Fc_AB_ and Fc_CD_ on opposite sides of. The SC has five immunoglobin domains (D1-D5), and each domain (D1-D5) are colored with different shades of cyan. The yellow asterisk marks the location of D1 CDR loops in unliganded and liganded structures and the orange asterisk indicated the location of the D1-D3 interface in the liganded (SIgA) structure. Color scheme and labeling approach are adapted from S. Kumar Bharathkar *et al.*, “The structures of secretory and dimeric immunoglobulin A,” *eLife*, vol. 9, p. e56098, Oct. 2020 (11).

Given critical roles in SIgA transport and function, the mechanisms of pIgR binding to pIgA have long been of interest(8),(15),(16),(17),(18). Earlier studies demonstrated that pIgR (and SC) D1 is both necessary and sufficient for binding to dIgA, but that D2-D5 further stabilize the SIgA complex(15),(16),(18),(19), and structural and biophysical studies have demonstrated that pIgR/SC domains undergo a massive conformational change upon dIgA binding (Fig.1) (8). The SIgA structures further revealed contacts between D1 and Fc_AB_ (the Fcα formed by HC-A and HC-B) and TpC and TpD, as well as D4-D5 contacts with Fc_CD_ (the Fcα formed by HC-C and HC-D)(11),(12),(13). Notably these interactions were accompanied by new inter-domain contacts between D1-D3 and D3-D4 not observed in the unliganded SC structure (Fig.S1). Furthermore, pIgR (and SC) binding to pIgA has long been appreciated to require the JC(4),(20),(21). SIgA structures suggest that the JC directly provides contacts for pIgR (or SC) D1 binding and indirectly provides the bent asymmetric conformation needed for other pIgR (or SC) domains to bind(11),(12),(13).

Taken together, pIgR binding to pIgA can be described as a multi-step process involving multiple interactions between pIgA and multiple SC domains (D1-D5), that undergo conformational changes to bind four IgA HCs and the JC, while also forming new inter-domain contacts within SC (Fig.1, Fig.S1); yet, aspects of this remarkably intricate mechanism remain unclear, such as the importance of inter-domain contacts within SC (D1-D5), and the role of JC and Tps in pIgR/SC binding. Building upon recent structural findings, we hypothesized that the conformational change-induced formation of pIgR/SC inter-domain contacts, together with binding to the C-terminus of JC is a critical step in SIgA assembly. To test this hypothesis, we carried out structure-based mutational analysis combined with Surface Plasmon Resonance (SPR) binding assays, that test the roles of pIgR/SC and JC residues in SIgA complex formation. Results signify that formation of the D1-D3 interface and D1 binding to the C-terminal residues of JC and Tps are pivotal steps necessary to achieve stable SIgA formation.

## MATERIALS AND METHODS

### Construct design and cloning

To make the mouse Fcα expression construct, the gene segment encoding the *mus musculus* IgA HC constant region (Uniprot P01878) C_H_2 and C_H_3 domains was codon optimized, fused to an N-terminal TPA signal sequence (residues MDAMKRGLCCVLLLCGAVFVSPAGA) and a C-terminal hexahistidine affinity tag and cloned into expression vector pD2610v1 (ATUM) using Electra cloning (ATUM), analogous to previously reported full-length IgA HC expression constructs(11). Constructs encoding the *mus musculus* JC (Uniprot P01592; native signal peptide) and *mus musculus* pIgR ectodomain residues 1-567 (Uniprot P01592; native signal peptide), were used as previously reported and served as templates for cloning truncated and/or mutant variants(11). SC and JC mutations were introduced into coding sequence fragments using overlap-extension PCR(22) with primers encoding the desired mutations (Integrated DNA Technologies; IDT) and Electra cloning sites; resulting products were cloned into expression vector pD2610v1 (ATUM) using Electra cloning (ATUM). SC-D5 cysteine mutations (C470A and C504A) were introduced into all SC expression constructs to prevent the formation of disulfide bonds with dIgA, as previously described(8). Constructs encoding the SC mD1-wt (residues 1-117) and mD1-mutant variants were generated by PCR and overlap extension PCR as described for other SC variants; coding sequence retained the native signal peptide and a C-terminal hexahistidine tag and were cloned into pD2610v1 using Electra cloning (ATUM). The DNA and amino acid sequences for expression construct coding regions are provided in a supplemental file.

### Protein expression and purification

Prior to expression, all expression construct plasmids were sequence verified (ACGT). Subsequently, constructs were transiently transfected (SC and D1 variants) or co-transfected (Fcα and JC) into Expi293F (Gibco: A14527) cells using the ExpiFectamine 293 Transfection kit (Gibco: A14525) and following previously described methods (11). Four to five days after transfection, supernatants were harvested and filtered through 0.22μm PES bottletop filters (Millipore Sigma). Proteins and complexes were purified from the cell supernatants using Nickel-NTA affinity resin (Qiagen: 30210), followed by Superose 6 (Cytiva) or Superdex 200 (Cytiva) size exclusion chromatography (SEC) on an AKTA FPLC (Cytiva). During and after SEC, purified molecules were maintained in TRIS-buffered saline (20mM Tris+ 150mM NaCl, pH=7.4). The SEC-eluted protein peaks corresponding to the correct retention volume based on the molecular weight standards (BioRad: 1511901) were collected and concentrated using Amicon Ultra centrifugal filters (Millipore Sigma) to stock concentrations between 2-4mg/ml for SC, 1.5-2mg/ml for D1, and 0.4-0.7mg/ml for dFcα constructs, and were then filtered using 0.22μm spin filters (Millipore Sigma) and utilized for SPR experiments.

### SPR experiments and data analysis

SPR experiments were carried out using a Sierra SPR-32 Pro (Bruker) operating at 25°C. Solutions containing 0.25μM-0.5μM dFcα-wt or dFcα mutant variant ligands in sodium acetate buffer (pH=4), were immobilized on lanes B1-8 and D1-8 of a High-Capacity Amine (HCA) 32-spot sensor (Bruker) using a standard amine coupling kit (Bruker). Lanes A1-8 and C1-8 were mock coupled using sodium acetate buffer pH=4 and served as references. The binding of soluble analytes, including mSC-wt, mSC-mutants, mD1-wt and mD1-mutants was tested using a two-fold (mSC and variants) or four-fold (mD1 and variants) dilution series starting at 2μM in HBS-EP+ running buffer (0.01 M HEPES, 0.15 M NaCl, 3 mM EDTA, 0.005% v/v Surfactant P20, pH 7.4). Analyte responses were measured on all spots and responses on lanes A and C (internal references) were subtracted to account for any non-specific binding. Kinetic experiments were carried out using a 25ul/min flow rate, 120s contact time, and 300s dissociation time. After every cycle the surface was regenerated using 2.5M MgCl2 three times, followed by a wash step with the running buffer (HBS-EP+) to achieve the best baseline for the next cycle. The SPR experiments testing mSC (wt and mutant variants) and mD1 (wt and mutant variants) analyte binding to dFcα-wt (data shown in Fig.2 and Fig.3) included five replicates. The SPR experiments testing mSC-wt/mD1-wt analytes binding to immobilized dFcα mutant variants (containing mutations in the JC) shown in Fig.4 were reproduced three or more times.

**Figure-2:**
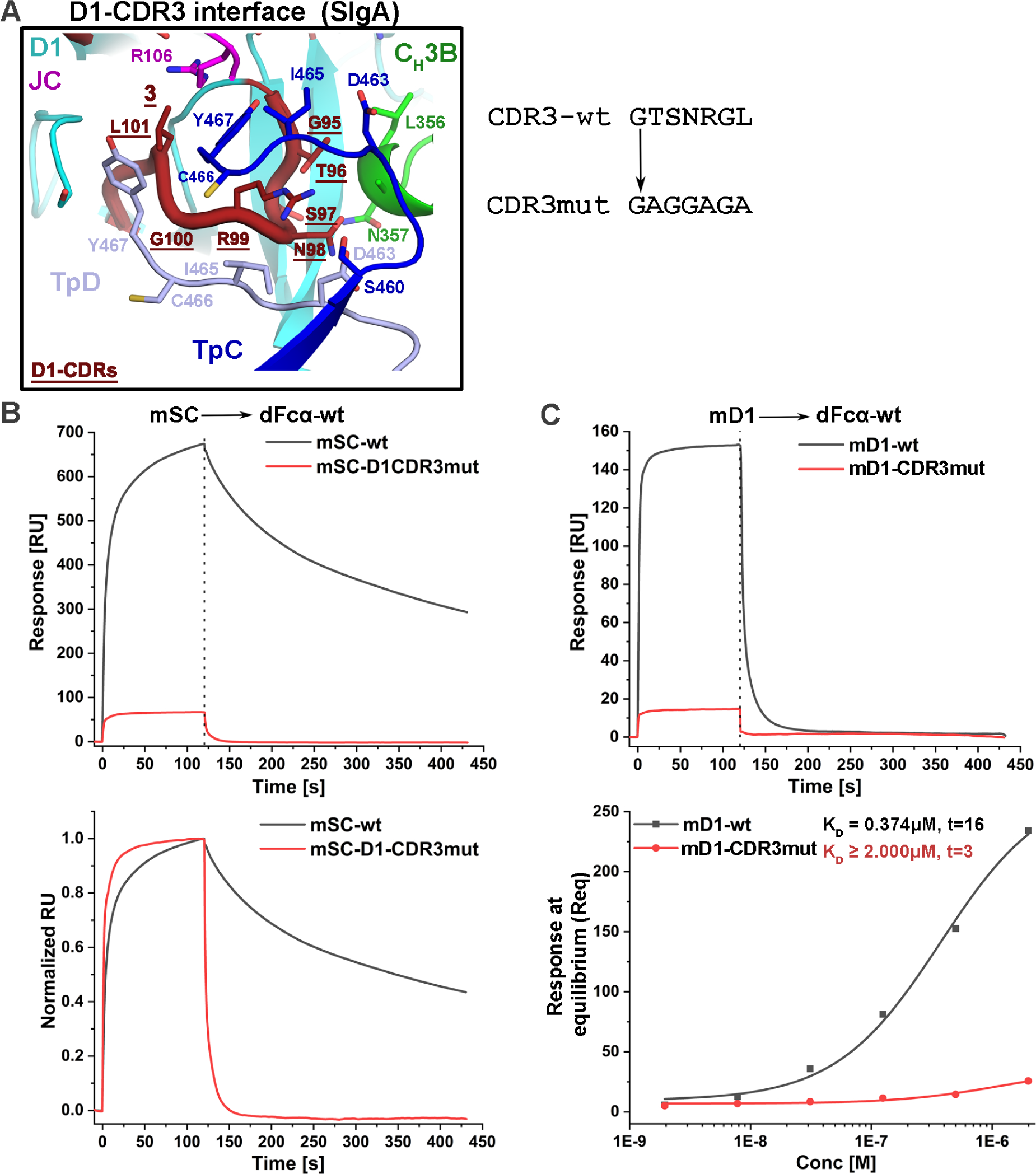
Contribution of mSC-D1-CDR3 towards binding dIgA. (A) Detailed view of mSC-D1-CDR3 and dIgA interactions found in mouse SIgA (PDB: 7JG2) (Fig.1, yellow asterisk); the structure is shown as a cartoon representation with interacting residues shown as sticks and labeled. D1-CDR3 residues that were mutated are underlined, and the unmutated and mutated sequences are shown on the right. (B) A representative SPR sensor-grams showing the response (top) and normalized response (bottom) of 0.5μM mSC-wt (black) and mSC-D1-CDR3mut (red) binding to dFcα-wt. (C) A representative SPR sensor-gram showing the response of 0.5μM of mD1-wt (black) and mD1-CDR3mut (red) binding to dFcα-wt (top) and the concentration dependent binding and steady state fit (bottom). The calculated K_D_ value for mD1-wt is indicated; the t-value associated with mD1-CDR3mut binding is t<10 indicating a poor fit and uncertainty in the calculated K_D_ value, which is estimated to be greater than 2 μM. Data from this figure are indicated by green inverted triangles in Fig.S5A. Results are consistent with 5 replicate experiments.

**Figure-3:**
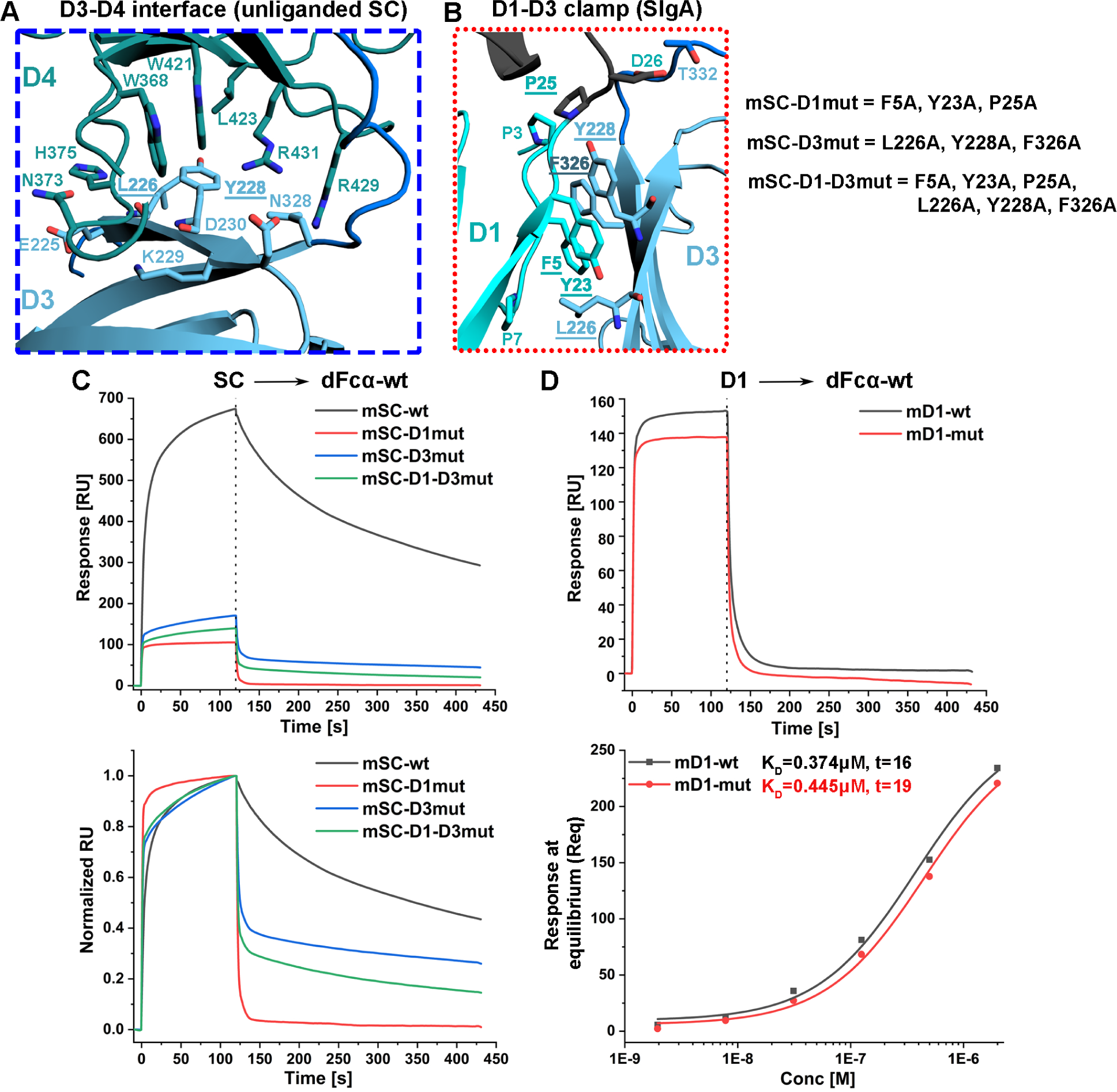
Role of SC-D1-D3 Clamp for the stabilization of SC and dIgA binding. (A) Detailed view of the unliganded D3-D4 interface from the homology model of mSC and the D1-D3 interface from the SIgA cryo-EM structure (Fig.1, orange asterisk) (PDB code 7JG2). In (A) and (B) structures are shown as a cartoon representation with interacting residues, shown as sticks and labeled. SC D1 and D3 residues that were mutated are underlined, and the unmutated and mutated sequences are shown on the right. (C) SPR sensor-grams showing the response (top) and normalized response (bottom) of 0.5μM mSC-wt (black) mSC-D1mut (red), mSC-D3mut (blue) and mSC-D1-D3mut (green) binding to dFcα-wt. (D) SPR sensor-grams showing the response of 0.5μM of mD1-wt (black) and mD1-mut binding to dFcα-wt (top) and the concentration dependent binding and steady state fit (bottom). The calculated K_D_ value for each analyte is given to the right of the name along with the t-value. Data from this figure are indicated by green inverted triangles in Fig.S5A. Results are consistent with 5 replicate experiments.

**Figure-4:**
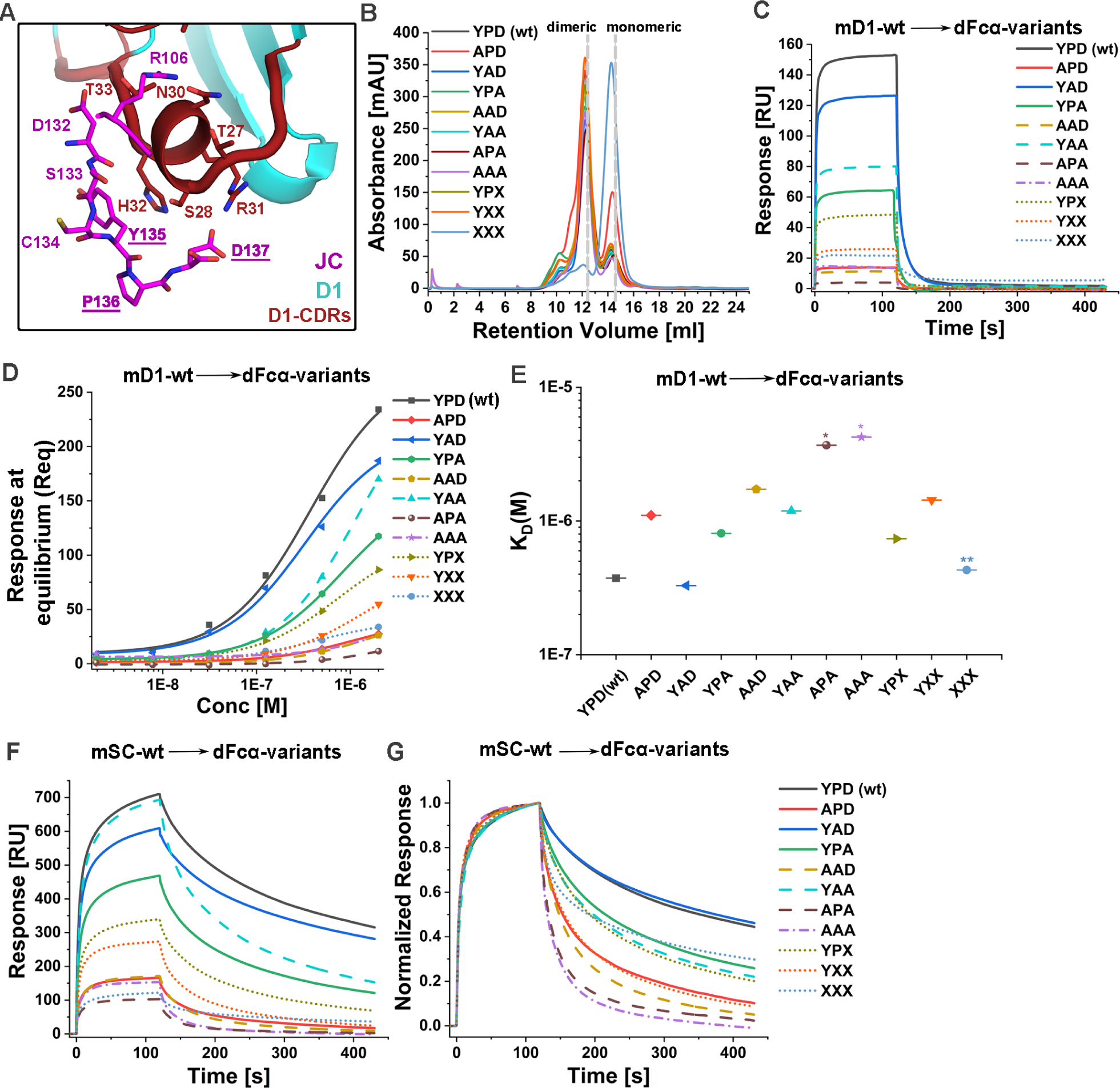
Role of the JC YPD motif in binding mSC-wt or mD1-wt. **(A)** The interaction of JC with SC-D1, and specifically D1-CDR1 shown as red loop with larger radius. The interacting residues are shown as sticks at this interface. Y135, P136 and D137 are underlined in JC, indicating that these residues were subjected to mutations, such as alanine substitutions or truncations. (B) The SEC chromatograms of the indicated dFcα-wt and dFcα mutant variants resulting from 25ml co-transfections and following Nickel-NTA purification; fractions corresponding to dimeric forms were used for SPR experiments. (C) SPR sensor-grams of 0.5μM mD1-wt binding to the indicated dFcαs with JC-YPD alanine mutations or truncations. (D) The steady state affinity analysis of mD1-wt binding to the indicated wt or mutant dFcα variants and (E) plot showing the resulting K_D_ values for the indicated variants. The * represents the data points for which the steady state fitting t-value was <10 and/or K_D_≥2μM indicating a poor fit; the affinity for data meeting these criteria are estimated K_D_≥ 2μM. ** represents a K_D_ value calculated from steady state analysis with t=11 and low maximal response indicating low confidence in the calculated value and estimated K_D_≥2μM. K_D_ values calculated from replicate experiments are summarized in FigS5B. (F) SPR sensor-grams of 0.5μM mSC-wt binding to different dFcα with the indicated JC-YPD alanine mutations or truncations. (G) SPR sensor-grams showing the normalized response of 0.5μM mSC-wt analytes binding to the indicated dFcα variants. Results are consistent with 3 or more replicate experiments.

Resulting SPR data were analyzed using the Bruker Sierra Analyzer software version R4. Briefly, all data were reference subtracted and buffer-subtracted and K_D_ values were calculated for the D1-only constructs using “steady state affinity” analysis program and 1:1 Langmuir + offset fitting model. The calculated K_D_ value for which the t-value is ≥10 is considered a reliable fit, where t-value is defined as K_D_/standard error of K_D_ by the Bruker Sierra Analyzer software version R4. Steady state equilibrium occurs when the quantity of molecules binding is equal to the quantity dissociating and the K_D_ is determined by fitting the response at equilibrium (Req) versus analyte concentration. The accuracy of curve fitting to the steady-state model is reduced when the K_D_ values are greater than half of the highest analyte concentration used (https://cdn.cytivalifesciences.com/api/public/content/digi-16798-pdf)(23). Given this, we have estimated that K_D_ values associated with t-values below 10 have an affinity K_D_ > 2μM (the maximum analyte concentration) to indicate that affinity has been reduced compared to unmutated controls, but the magnitude of reduction is uncertain. The degree of uncertainty is evident by comparison of K_D_ values calculated from replicate experiments (Fig.S5), which exhibit a greater range for mutant variants with high K_D_ values (e.g. above 2 μM) or t <10 compared to unmutated controls. Despite this variation, the overall result (a reduction in binding associated with mutations) is consistent across all replicates. Raw sensor-grams (RU vs Time) for all data and the steady state affinity data for mD1 variants (Reqvs Conc), were exported and re-plotted using OriginPro 2020 graphing and analysis software (OriginLab: https://www.originlab.com/). Normalization of the respective sensor-grams for concentration matched samples were carried out in OriginPro 2020, by dividing RU by the maximum Response of the respective analyte. Figures were assembled and in Photoshop (Adobe) or Inkscape (https://inkscape.org/).

## RESULTS

To investigate the molecular mechanisms underlying SIgA assembly we analyzed existing SC(8), dIgA(11) and SIgA(11),(12),(13) structures and identified SC and JC residues with potential to mediate the pIgR binding mechanism, for example, residues that may promote SC conformational changes and/or stabilize SIgA interfaces. Subsequently, we mutated target residues in recombinantly expressed full-length SC and D1-only variants and tested binding to immobilized recombinant dIgA lacking the Fabs (dFcα) using SPR binding assays; we also tested unmutated SC and D1-only binding to dFcα containing JC mutant variants. SC and D1 analytes provide an established, soluble model for pIgR binding to dIgA(8). Consistent with the ligand-binding-induced conformational change observed for full-length SC, published SC-dIgA binding studies do not fit single state kinetic models, and thus similar to prior studies we followed the approach of qualitatively monitoring differences in the SPR binding profiles to identify differences among protein variants(8),(24), while also calculating equilibrium binding affinities for variants containing only D1. All experiments utilized mouse SC, Fcα and JC sequences.

### D1-CDR3 plays a crucial role in the initiation of SC binding

SC D1-D5 Ig-like domains each contain up to three loops (1–3) that are structurally similar to antibody complementarity determining regions (CDRs)(8). SIgA structures revealed residues in SC D1, including those in CDRs 1-3, mediating contacts with all four HCs and JC (Fig.S1C and Fig.S1F). However, the unliganded SC crystal structure revealed that a subset of these residues, including those in D1-CDR1 and CDR2, interact with D4 and D5 leaving them sterically occluded and likely unavailable to initiate binding between SC-D1 and dIgA, while D1-CDR3 is solvent exposed and does not participate in any interdomain interactions (Fig.S1C and Fig.S1F). This observation led us to hypothesize that D1-CDR3, which is solvent exposed and sterically accessible in unliganded SC, initiates pIgR binding to dIgA.

In SIgA structures, SC-D1-CDR3 residues contact HC-B, TpC and TpD though hydrogen bonds and ionic interactions, and additionally, lie within 3.5-6Å of JC, where they may mediate Van-der Waals interactions (Fig.2A). To explore the hypothesis that SC-D1-CDR3 is crucial for SC and dIgA binding, we generated SC expression constructs, in which the SC-D1-CDR3 residues were mutated to Glycine and Alanine (CDR3-wt: GTSNRGL to CDR3-mut: GAGGAGA), limiting the possible side chain interactions with dIgA, while preserving the length of the loop (Fig.2A). The first construct incorporated this mutation into full-length SC (mSC-D1-CDR3mut) and the second incorporated it into an expression construct encoding only SC-D1 (mD1-CDR3mut). We expressed and purified variants along with unmutated controls, mSC-wt and mD1-wt, and tested their binding to dFcα-wt (Fig.2 and Fig.S2).

The mSC-D1-CDR3mut SPR sensor-grams revealed markedly lower response units (RU) compared to concentration-matched mSC-wt responses, with an overall maximum measured RU of mSC-D1-CDR3mut that was ∼9.9% of the mSC-wt RU for the 0.5μM analyte sample (Fig.2B and Fig.S2A). Furthermore, overlay of normalized responses revealed more rapid association and dissociation for mSC-D1-CDR3mut compared to mSC-wt (Fig.2B). Similarly, the mD1-CDR3mut SPR sensograms revealed markedly lower response, ∼9.6% of the concentration-matched mD1-wt response (Fig.2C, Fig.S2B). Reduction in mD1-CDR3mut responses over the tested concentrations limited the accuracy of steady-state analysis; however, suggest an overall reduction in affinity from K_D_=0.374μM for mD1-wt to K_D_ ≥2μM for mD1-CDR3mut (Fig.2C, Fig. S5A). In contrast to comparisons between mSC-D1-CDR3mut and concentration matched mSC-wt, the responses revealed similar association and dissociation phases for mD1-CDR3mut and mD1-wt. Taken together, these data indicate that mutations in D1-CDR3 reduce D1 binding affinity in the context of SC and isolated D1, but alter the binding kinetics only in the context of SC. These results point toward a central role for D1-CDR3 in promoting D1-dIgA interactions, but also a role in promoting changes in other SC domains upon dIgA binding, as evidenced by differences in binding kinetics for mSC-wt and mSC-D1-CDR3mut. These differences may result from conformational changes that expose more of D1 for binding and/or the binding of other SC domains.

### The formation of the SC D1-D3 clamp is crucial for stabilization of SC-binding

In unliganded human SC, D1 forms extensive interfaces with SC domains D2, D4 and D5, yet shares minimal contacts with D3 (no contacts in mouse SC, one contact in human SC)(8). However, in the SIgA structure, D1 contacts only D3 through an extensive hydrophobic interface described as a “D1-D3 clamp” (Fig.3B)(11),(12),(13). The hydrophobic interface involves D1 residues F5, P7, Y23, P25 and D3 residues L226, Y228, F326. Notably, D3 L226 and Y228 contact SC D4 in the unliganded SC structure, implying that D3 swaps interfaces between D4 (unliganded) and D1 (liganded) during SC binding in what we hypothesize is a dominant step in SC’s conformational change associated with SIgA formation (Fig.3A and Fig.S1D, S1E).

To investigate the significance of the D1-D3 clamp, we generated three mutated SC variants to minimize the hydrophobic interface: (1) mSC-D1mut, which includes D1 F5A, Y23A, and P25A mutations, (2) mSC-D3mut, which includes D3 L226A, Y228A, F326A mutations, and (3) mSC-D1-D3mut, which includes D1 F5A, Y23A, P25A and D3 L226A, Y228A, F326A mutations (Fig.3B) and tested their binding to mouse dFcα-wt. Compared to mSC-wt, SPR sensor-grams revealed lower overall RUs and lower maximum measured RU for the binding of concentration-matched mSC-D1mut (15.7% mSC-wt RU), mSC-D3mut (25.3% mSC-wt RU), and mSC-D1-D3mut (20.7% mSC-wt RU) (Fig.3C and Fig.S3), indicative of weaker binding. Qualitatively, the reduction in RU appeared similar for all three mutant variants, with maximum measured RU values being between 15-26% of mSC-wt for 0.5μM analytes; the combination of mutations in both D1 and D3 did not appear to produce an additive effect, indicating that disruption of interface residues in either D1 or D3 was sufficient to disrupt the interface.

Overlay of concentration matched, normalized responses for mSC-wt and mSC-mutant variants also revealed differences in the binding kinetics, with mSC-D1mut exhibiting faster association and faster dissociation phases compared to mSC-wt (Fig.3C). Responses of mSC-D1mut were similar to those of mD1-wt (Fig.3C, Fig.3D), which may indicate that only D1 (and not D4 and D5) is binding. However, mSC-D3mut and mSC-D1-D3mut exhibited association phases similar to mSC-wt, but faster dissociation phases compared to mSC-wt. Compared to mSC-D1mut, mSC-D3mut and mSC-D1-D3mut exhibited moderate differences in the dissociation phase compared to mSC-D1 mut, suggesting that mutation of the D3 interface may impact binding differently than mutation of the D1 interface (Fig.3C and Fig.S3).

To explore the possibility that mutation of D1 F5A, Y23A, P25A influenced D1 binding independent of the other SC domains we generated mD1-mut, which contained D1 F5A, Y23A and P25A mutations, and compared its binding affinity to mD1-wt (Fig.3D and Fig.S3). The mD1-mut binding affinity, K_D_=0.445μM, was comparable to the mD1-wt binding affinity, K_D_=0.374μM, indicating that D1 interactions with dIgA were not disrupted (Fig.3D and Fig.S5A). Thus, in the context of full-length SC, the mutations F5A, Y23A and P25A appear to reduce binding to dIgA indirectly, presumably through disruption of the D1-D3 clamp. Taken together, results imply that D1-D3 interface formation contributes to stable SIgA assembly but is not necessary for SC (or D1) binding. As noted, the kinetics of the three D1-D3 interface mutants closely resemble that of isolated SC D1 (Fig.3 and Fig.S3) and thus it is plausible that disruption of the D1-D3 clamp prevents subsequent stages of SC binding (e.g., part of the conformation change and D4-D5 binding to dIgA) resulting in only D1-dIgA contacts. Alternatively, D1-D3 clamp disruption may markedly decrease the efficiency of SC D4-D5 binding and/or the stability of the resulting SIgA.

### JC C-terminal residues mediate pIgR binding to dIgA

The structures of human and mouse SIgA reveal similar conformations for SC, JC and similar contacts between chains. Among these interactions, SC-D1 contacts with JC are conserved, and contribute to 63% of the total SC-JC interface. Of the three SC-D1 CDRs, JC interfaces primarily with SC-D1-CDR1, forming hydrogen bonds and ionic interactions between JC residues R106, D107, D132, Y135 and D137 and SC-D1-CDR1 residues S28, D30, R31, H32 and T33 (Fig.4A). JC Y135 and D137 are part of the conserved C-terminal *YPD* motif that follows JC wing-2; JC-YPD are disordered in the dIgA structure(11) and therefore we hypothesized that these residues play a dominant role in pIgR binding, rather than dIgA assembly. Furthermore, in unliganded SC, D1-CDR1 R31 and H32 form contacts with D4-D5(Fig.S1C)(8) and thus, JC YPD may function to displace the D1-D4D5 interface and/or promote D1-JC binding while freeing D4-D5 to bind the Fc_CD_. To investigate these possibilities, we created ten JC variants with mutations or truncations in YPD.

Initially we tested the effect of YPD mutations or truncations on dFcα assembly by co-expressing each JC variant with Fcα and found that all could be incorporated into dFcα, indicating that mutations did not inhibit dFcα assembly. In most cases, the relative percentage of dFcα and monomeric Fcα, which is found in lower abundance in all dFcα preparations, was similar for preparations containing unmutated or mutated JC. However, truncation of YPD resulted in higher percentages of monomeric Fcα and lower dFcα yield, suggesting that the efficiency of JC incorporation, and therefore dIgA assembly, may rely in-part on the presence of JC C-terminal residues (Fig.4B). Accordingly, we used mutated dFcα variants as immobilized ligands in mD1-wt and mSC-wt SPR binding assays.

Initially, we tested binding of mD1-wt to dFcα-wt and dFcα-mutant variants having single (dFcα-APD, dFcα-YAD, dFcα-YPA), double (dFcα-AAD, dFcα-ADA, dFcα-YAA) or triple (dFcα-AAA) alanine substitutions in JC YPD and calculated the binding affinity for each variant. Results revealed binding affinities of K_D_=0.374μM, K_D_ = 1.104μM, K_D_ = 0.329μM and K_D_ = 0.810μM for dFcα-wt and single alanine variants dFcα-APD, dFcα-YAD, and dFcα-YPA, respectively (Fig.4C, Fig.4D, Fig.4E, Fig.S5B). Binding affinities for double mutants dFcα-AAD, and dFcα-YAA were K_D_ =1.728μM and K_D_ =1.191 μM, respectively. mD1-wt binding to dFcα-APA exhibited the lowest responses among double alanine substitutions, reducing the accuracy of steady-state analysis and suggesting a reduction in affinity to K_D_ ≥2μM (Fig.4C, Fig.4D, Fig.4E, Fig.S4, Fig.S5B). Collectively, these results demonstrate a marked reduction in binding affinity when either Y135 or D137 is mutated and suggest a further reduction in binding affinity when both Y135 and D137 are mutated to alanine. The P136A mutation resulted in comparatively minor changes in binding on its own; it may further reduce binding when combined with Y135A or D137A mutations. The mD1-wt binding to triple mutant dFcα-AAA also exhibited low responses, limiting the accuracy of steady-state analysis and calculated affinity, which we estimated to be K_D_ ≥2μM. Results further indicate that the substitution of YPD residues has a pronounced effect on binding (Fig.4C, Fig.4D, Fig.4E, Fig.S4, Fig.S5B).

We also tested the mSC-wt binding to all dFcα YPD-alanine substitutions. Although resulting SPR responses do not fit a single-state kinetic model or reach equilibrium during the experiment, differences in dissociation phases for concentration matched, normalized responses were apparent when comparing mSC-wt binding to dFcα-wt and dFcα YPD-alanine substitutions (Fig.4F, Fig.4G). Specifically, we observed that all mutations except P136A altered dissociation. Consistent with experiments that tested D1 binding, single mutation Y135A exhibited the most pronounced effect, followed closely by D137A; the combination of Y135A and D137A further altered the dissociation phase (more rapid dissociation) and the triple mutant containing Y135A, P136A, and D137A exhibited the most significant change in dissociation compared to unmutated control, dFcα-wt. Qualitatively, we also observed subtle differences in association and overall reduction in maximum measured RU values for mSC-wt binding to the majority of dFcα YPD-alanine substitutions (Fig.4F and Fig.S4).

Taken together YPD-alanine substitutions reveal a central role for JC-YPD in mediating D1 binding to dIgA. Y135 and D137 appear to play prominent and unique roles since individual mutations in these residues reduced binding affinity whereas mutation of P136 alone did not. Both Y135 and D137 appear to contribute uniquely since dFcα YPD-alanine substitutions that combined them exhibited further reductions in binding. While a more minor role for P136 was observed, its mutation to alanine did reduce binding affinity, most notably when combined with the Y135A and/or D137A mutations. Notably, none of the dFcα YPD-alanine substitutions completely abrogated mD1-wt or mSC-wt binding indicating that either YPD main-chain interactions and/ or other JC-SC interfaces are sufficient for SIgA formation in our experiments. To investigate the role of JC length and YPD main chain atoms on interactions with SC, we tested SC binding to JC deletion mutants, in which D137 (dFcα-YPX), P136 and D137 (dFcα-YXX), or Y135, P136 and D137 (dFcα-XXX) were deleted. Initially, we tested mD1-wt binding to dFcα-wt and dFcα YPD deletions, dFcα-YPX, dFcα-YXX and dFcα-XXX and calculated associated binding affinities. Results revealed binding affinities of K_D_= 0.737μM, and K_D_= 1.434μM for dFcα-YPX, dFcα-YXX, respectively. The mD1-wt binding to dFcα-XXX exhibited the lowest responses among deletion mutants, reducing the accuracy of steady-state analysis and indicating an overall reduction in affinity estimated at K_D_ ≥2μM (Fig.4C, Fig.4D, Fig.4E, Fig.S5B). Collectively, data indicate a reduction in binding upon the deletion of one or more JC C-terminal residues compared to dFcα-wt K_D_=0.374μM, indicating that a change in JC length can impact D1 binding. We also tested the mSC-wt binding to all dFcα YPD-deletion variants and similar to dFcα YPD alanine substitutions, observed differences in the dissociation phases for concentration matched, normalized responses when comparing mSC-wt binding to dFcα-wt, dFcα-YPX, dFcα-YXX and dFcα-XXX (Fig.4F, Fig. 4G). While we observed more rapid dissociation for mSC-wt binding to dFcα-YPX and dFcα-YXX (compared to dFcα-wt), binding to dFcα-XXX exhibited dissociation more similar to dFcα-wt, albeit still altered. Taken together, these results further support a role for JC YPD in SIgA formation and suggest that both sidechain identity and the presence of mainchain atoms support the interaction.

## DISCUSSION

The pIgR binding to dIgA is a critical step in SIgA formation, its transcytosis to mucosal secretions, and downstream effector functions(25),(26). Early studies conducted both *in vitro*, using SC (15) (16), and in cells expressing pIgR (19),(18) demonstrated that D1 is necessary and sufficient for dIgA binding, identified involvement of all three D1 CDR loops and identified covalent interactions between D5 and dIgA(27),(28),(29). These reports, together with the observation that pIgR is tethered to the cell surface via a long (∼30 residues in humans) linker that is predicted to be flexible, led to subsequent structural and biophysical studies using SC as a model for pIgR (8). Results unveiled an intricate, multi-step process, in which domains D1-D5 undergo conformational rearrangements that uncover D1 CDR1 and CDR2 residues and reposition D4 and D5 to facilitate dIgA binding (8). This model is consistent with recently published cryoEM structures which reveal D1 CDR residues contacting all four HCs and the JC and D4 and D5 bind a single HC (8), (11),(12),(13). Here we characterized the contributions of specific residues in the D1 CDR3, the D1-D3 clamp and the JC C-terminal YPD residues, which provide new details on how SIgA assembles.

In mouse SIgA, the majority of D1 CDR contacts involve residues in CDR1, which contains a conserved helical turn and has long been thought to play a pivotal role in the SIgA assembly process (15), (11). Our data also point toward a central role for D1-CDR3 in this process; D1-CDR3 residues account for just 34.3% of D1’s total buried surface, yet in our studies mutation of these residues reduced D1 binding affinity and altered SC association and dissociation phases. This is consistent with early studies using rabbit pIgR, which demonstrated that substitution of D1 CDR loops with sequences from other SC domains limited dIgA binding (18). As noted, D1-CDR3 residues, unlike D1-CDR1 and D1-CDR2 residues, are fully exposed in the unliganded SC structure(8) and thus, our results are consistent with the model in which exposed D1-CDR3 residues are able to initiate binding by contacting Tps and JC residues, promoting D1-CDR1 and D1-CDR2 contact with dIgA, and likely initiating the conformational change. CDR3 residues also appear to contribute considerable stability to SC interactions with pIgA in the fully assembled structure and interface with at least four different chains. In the mouse SIgA structure(11), SC-D1-CDR3 residues participate in a network of hydrogen bonds and ionic interactions with HC-B, TpC, and TpD as well as putative Van-der Waals interactions with the JC. Furthermore, these Tps (specifically TpD) and JC residues appear to be flexible (disordered) in the dIgA structure (11) and we speculate that binding to CDR3 helps stabilize their positions and following the conformational change and promote additional interactions with D1 (e.g. CDR1 and CDR2).

D1-CDR3 may also promote binding to higher order pIgA and pIgM. Our own inspection of the human dimeric (PDB: 6UE7 and 6LX3)(12),(13), tetrameric (PDB: 6UE8 and 6UE9), and pentameric (PDB:6UEA) SIgA core structures(12), reveals D1-CDR3 contacting one Tp (the others are not resolved) in dimeric SIgA and contacting two Tps in tetrameric and pentameric SIgA; in pentameric SIgA, a third Tp contacts D1 outside the CDR regions (11),(12),(13). These observations are consistent with earlier reports that SC bound tetrameric IgA2m2 with better affinity than dimeric IgA1 and dimeric IgA2m1(5) and suggest D1-CDR3 may support high affinity binding to higher order pIgA. Our inspection of the related SIgM core structure (PDB: 6KXS and 7K0C)(30),(31) indicates that SC D1 forms additional contacts with the Tps in pentameric SIgM (similar to pentameric SIgA and in contrast to dimeric SIgA) and thus evolution of D1 may have been driven by interactions with higher order polymers of IgM and IgA and their respective Tps.

The structures of SIgA (and SIgM) revealed that the pIgR’s ligand-induced conformational change is associated with the formation of the D1-D3 clamp and data presented here revealed that disruption of the D1-D3 clamp markedly reduces dIgA binding. Structurally, the unliganded SC D3-D4 interface must be disrupted for D1-D3 interface to form. This implies that either the D1-D3 interface forms first and thereby frees D4-D5 to bind dIgA, or that D4-D5 first binds to dIgA and thereby frees D3 to bind D1. Indeed, SC variants with mutated clamp residues exhibited SPR binding profiles that are similar to that of D1 only, which may indicate that in the absence of the clamp, D4 and D5 are not able to bind efficiently. While this could result from failure of the clamp to form and/or from the failure of the unliganded SC D3-D4 interface to break; it is notable that among D1-D3 interface mutants, mSC-D1mut is the weakest binder. The mSC-D1mut lacks the L226A and Y228A mutations found in mSC-D3mut and mSC-D1-D3mut, which are expected to disrupt the D3-D4 interface in unliganded SC. Thus, it appears plausible that D1-D3 clamp formation frees D4 and D5 and positions them optimally to bind dIgA. Regardless, it is clear that both disruption of unliganded SC D3-D4 contacts and formation of the liganded D1-D3 clamp and are crucial for the formation of a stable SIgA complex. We also speculate that the formation of the D1-D3 clamp, which effectively locks D1 (bound to one IgA) to D4-D5 (bound to the other IgA), may impact SIgA functions. Comparison of mouse dIgA and SIgA structures identified conformational differences primarily manifesting as different angles tilt between the two IgA monomers (30 degrees for SIgA and 19 degrees for dIgA). These differences imply that SC influences the global conformation of the SIgA core, a result which computational modeling predicts may influence the positions of Fabs and therefore, antigen binding (11).

The JC has long been known to be necessary for dIgA assembly in mammals and for pIgR binding to dIgA; yet gaps in our understanding of how it mediates both processes remain (32). The mouse dIgA structure revealed numerous contacts between JC and two copies of mIgA; however, the conserved C-terminal JC YPD motif was disordered (11), hinting at a limited role in dIgA assembly. Our data demonstrate that dIgA assembly can tolerate mutations in JC YPD, and thus, its identity is not necessary for dIgA assembly. A limited role for the JC C-terminus in dIgA assembly is consistent with earlier studies, in which deletion of the 25 or more residues from the C-terminus or substitution with JC from other species (which encode different C-terminal sequences) did not completely abrogate polymeric IgA assembly but in some cases limited SC binding (17)(21). However, our data indicates the presence of three C-terminal residues in the YPD location may contribute to the dIgA assembly process since deletion of all three residues resulted in markedly lower levels of dFcα expression. Additionally, the mouse SIgA structure revealed an ordered YPD motif contacting SC-D1 (11) and we find that mutations of these residues, particularly Y and D, markedly impact SC binding. Yet, mutation or deletion of JC YPD doesn’t completely inhibit SC binding *in vitro*. Thus, other contacts between the JC and SC and/or dIgA appear to be sufficient for binding in our experiments. While these contacts may include other JC-SC interfaces, it may also be that SC-SC and/or SC-HC contacts help compensate for JC mutations and that those contacts are dependent on the overall bent dIgA conformation, conferred by largely by the JC (but not directly by the YPD). Regardless, our data support the notion that direct contacts with the JC C-terminal YPD motif are an inherent part of the pIgR-dIgA binding mechanism.

Notably, none of our SC or JC mutant variants completely abrogated SC-dIgA binding. This is valuable for determining the relatively contributions of mutated residues; however, we speculate that the reductions in binding affinity and apparent changes in kinetics that we observe could translate to severe disruption of pIgR-dependent transcytosis and or reduced SIgA stability in the mucosa. For example, mutations in the D1-D3 clamps and/or mutations disrupting interfaces may increase the exposure and/or flexibility of SC and/or dIgA motifs making them more susceptible to proteolysis. They may also limit subsequent stages of pIgR binding to dIgA. For example, mD5 C470 (C468 in human) can form a disulfide bond with HC C306 (C311 in human IgA) that stabilizes SIgA after SC D5 has bound dIgA (11),(12),(13),(27),(28). As noted in our results, mutations in SC D1 and/or D3 resulted in kinetic profiles similar to D1 only variants, and thus these mutations may limit D4/D5 binding and subsequent disulfide bond formation; however, in our SPR experiments (and previously reported binding studies (8),(24)) SC C470 and C504 were mutated to alanine to prevent irreversible binding to dIgA surfaces.

As noted above, decades of research have utilized SC as a model to describe pIgR interactions with dIgA (or IgM) and specific domain and residue contributions to binding are largely in agreement with molecular contacts observed in unliganded SC and SIgA structures, supporting the notion that the SIgA structure is essentially superimposable with the pIgR-dIgA complex tethered to cell membrane. However, we note that it is possible that intermediate stages of pIgR binding occur on different timescales *in vivo* than (SC binding) *in vitro* and that that during transcytosis, copies of pIgR-dIgA might exist in different binding stages (e.g. D1 bound but D4 and D5 unbound). However, upon D4/D5 binding, and the formation of the disulfide that links D5 to dIgA, the binding reaction would be complete and essentially irreversible under oxidizing conditions (e.g. endosomal or extracellular). It also remains unknown exactly when pIgR is proteolytically cleaved off the cell membrane; it is possible that the final stages (e.g. binding of D5) can occur during or following cleavage.

## AUTHOR CONTRIBUTIONS

The study was conceived by B.M.S and S.K.B.; experiments were conducted by S.K.B; Authors contributed to data analysis and manuscript writing.

## ACKNOWLEDGEMENTS

We thank Paul Ritter and Hniang Khamh for help with SPR training and troubleshooting, Sarah Leonard for technical assistance, and members of the Stadtmueller Lab for insightful discussions regarding this work.

## Supplementary Information

**Supplemental Figure-1:**
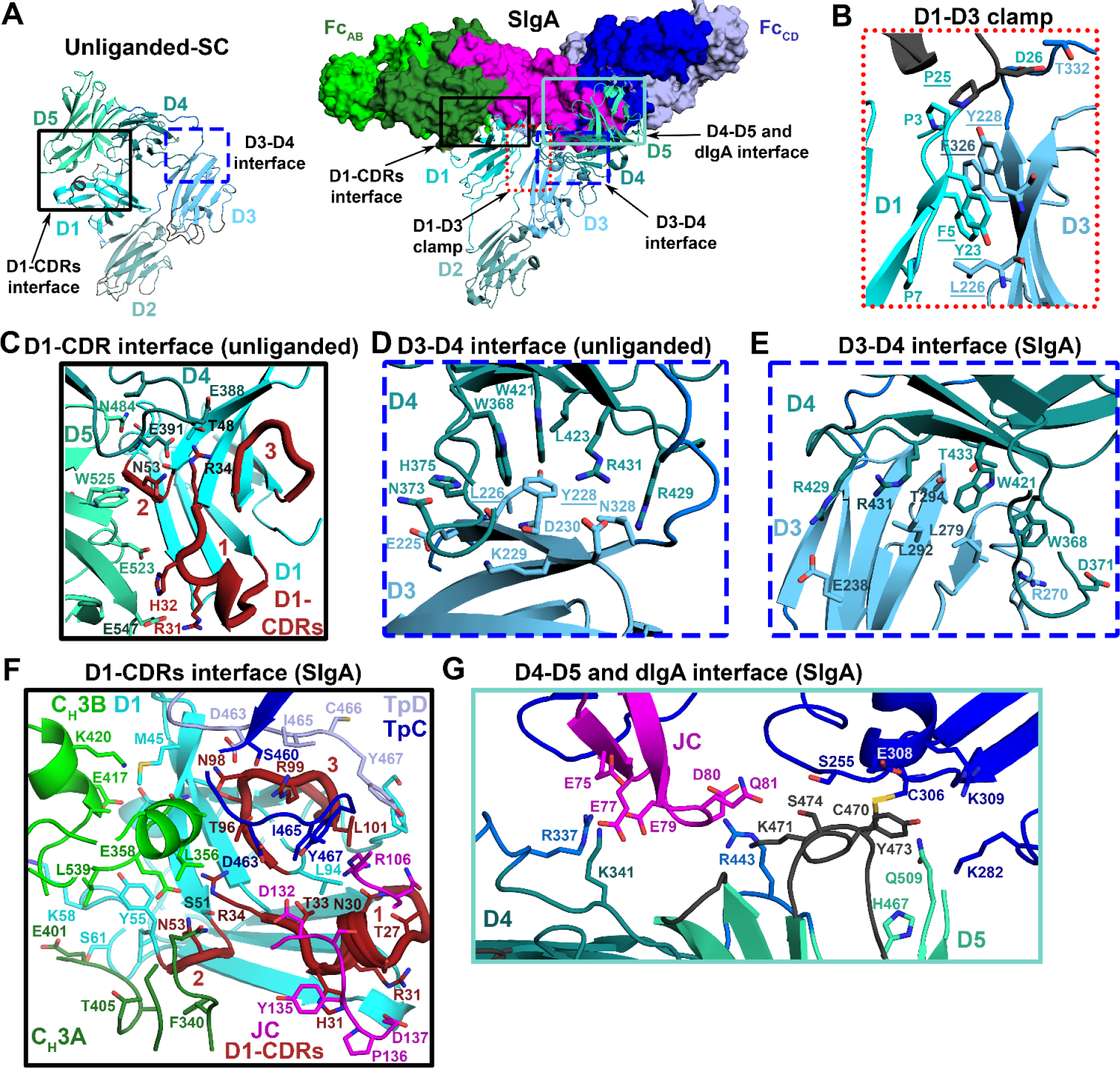
Different interfaces of SC-domains in unliganded and liganded structures. (A) Homology model of unliganded mouse SC based on the unliganded human SC structure (PDB: 5D4K) and the mouse SIgA Cryo-EM structure (PDB: 7JG2). The SC is shown as a cartoon, with its five domains (D1-D5) colored different shades of cyan. In SIgA, dIgA components are shown as a molecular surface and colored as Fig. 1. The boxed regions are enlarged in B-G and detail residues engaged in molecular interactions for the (B) liganded SC D1-D3 clamp in SIgA, (C) unliganded SC D1 and D4-D5 interface, (D) unliganded SC D3 and D4 interface, (E) liganded D3 and D4 interface in SIgA, (F) liganded D1 and dIgA interface (G) the liganded D4-D5 and dIgA interface. In all panels, interfacing residues are shown as sticks. The D1-CDRs are shown as thicker loops denoted as 1, 2 and 3 corresponding to CDR1, CDR2 and CDR3 respectively, and they are colored brick red.

**Supplemental Figure-2:**
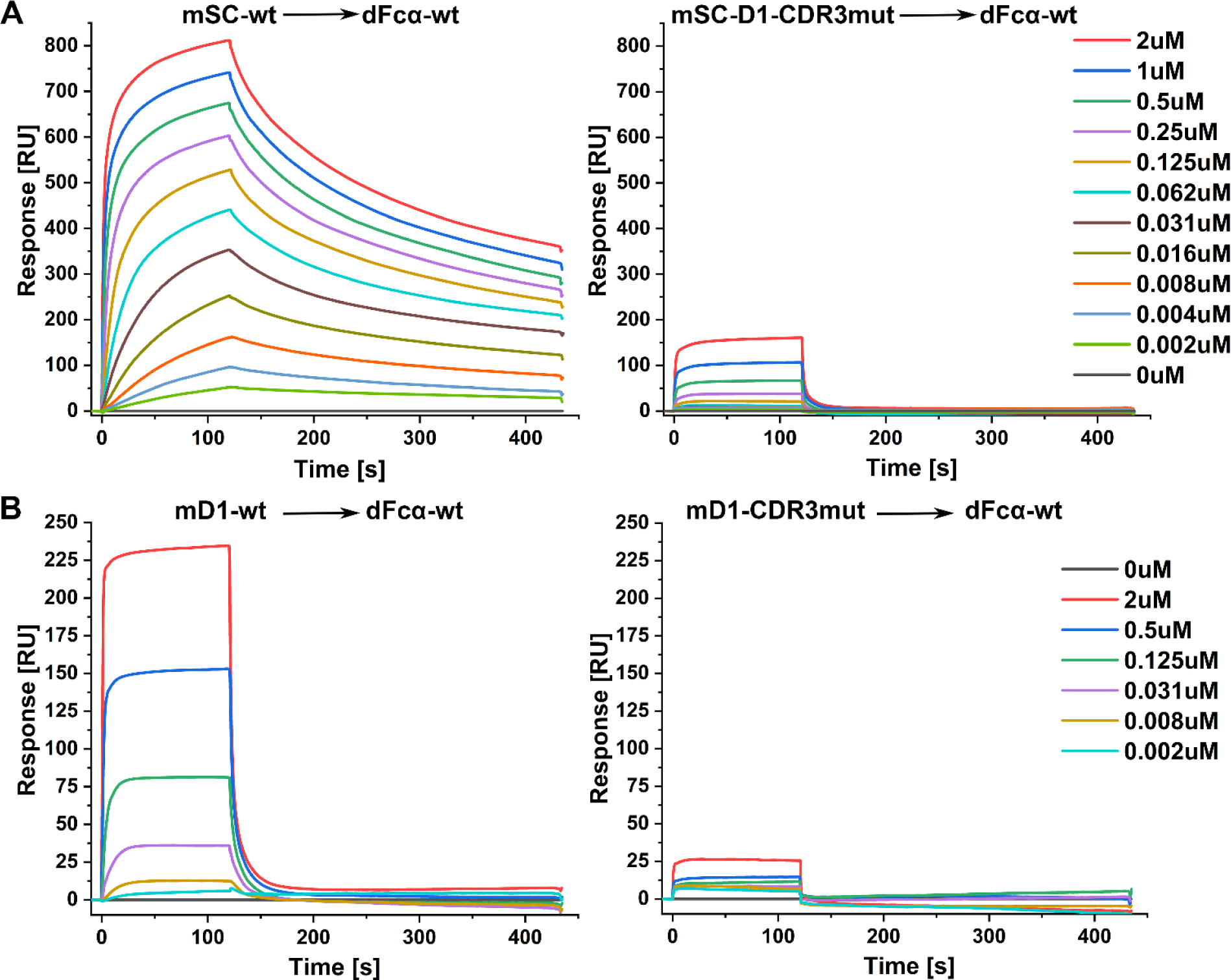
mSC-wt and mSC-D1-CDR3mut binding to dFc*α*-wt at various concentrations. (A) SPR sensorgrams showing the responses for the entire dilution series of mSC-wt and mSC-D1-CDR3mut binding to dFcα-wt, shown with the same scale. (B) SPR sensorgrams showing the responses for the entire dilution series of mD1-wt and mD1-CDR3mut binding to dFcα-wt. The responses for each analyte dilution are colored according to the keys (right), shown with the same scale. Results were consistent between 5 replicate experiments.

**Supplemental Figure-3:**
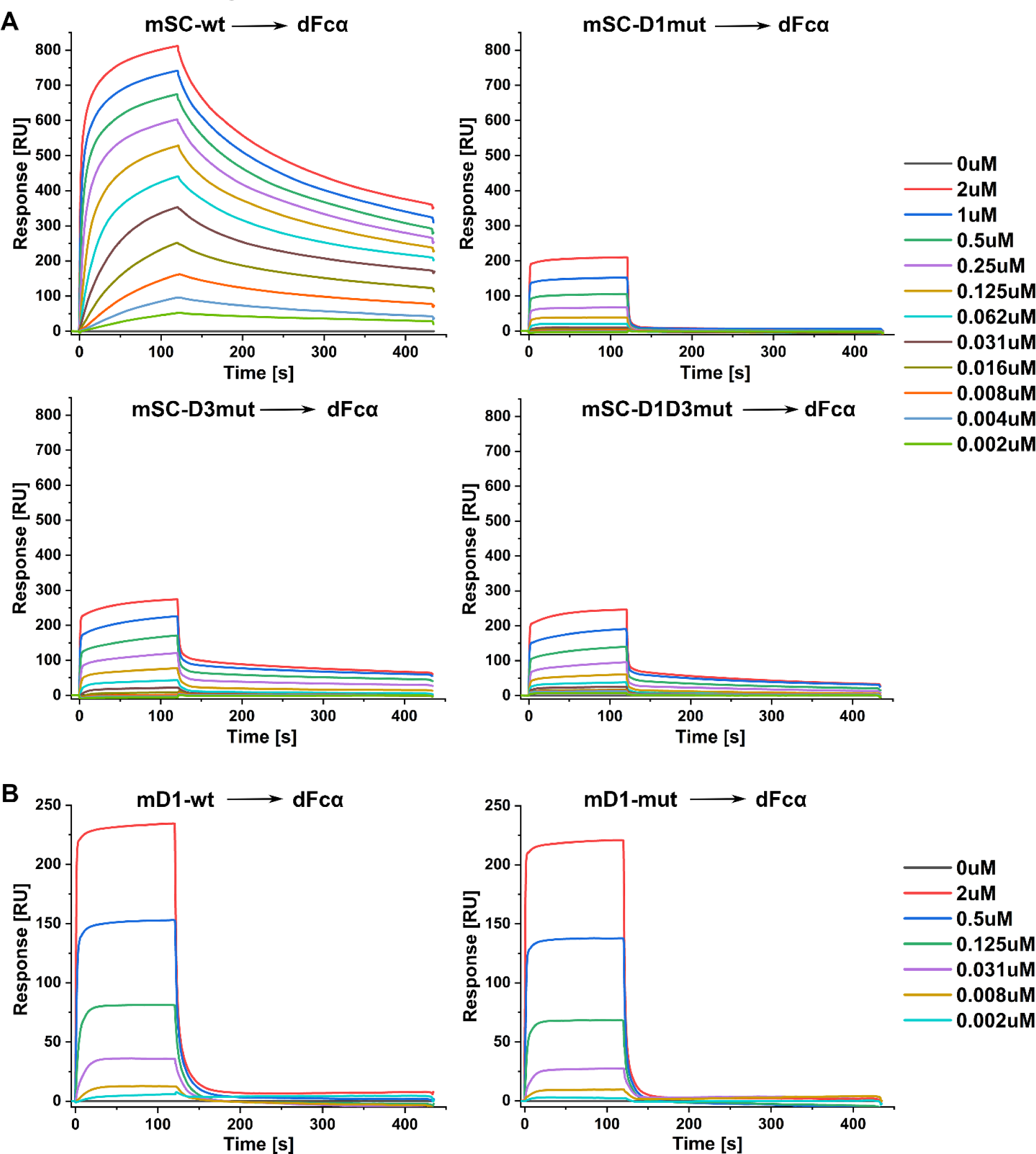
Role of D1-D3 in binding stabilization and conformation change. (A) SPR sensorgrams showing the responses for the entire dilution series of mSC-wt, mSC-D1mut, mSC-D3mut and mSC-D1-D3mut, binding to dFcα-wt (2-fold dilution series starting from 2uM). All shown in same scale for comparison. (B) SPR sensorgrams showing the responses for the entire dilution series of mD1-wt, and mD1mut, binding to dFcα-wt (4-fold dilution series from 2uM). The responses for each analyte dilution are colored according to the keys (right). Results were consistent among 5 replicate experiments.

**Supplemental Figure-4:**
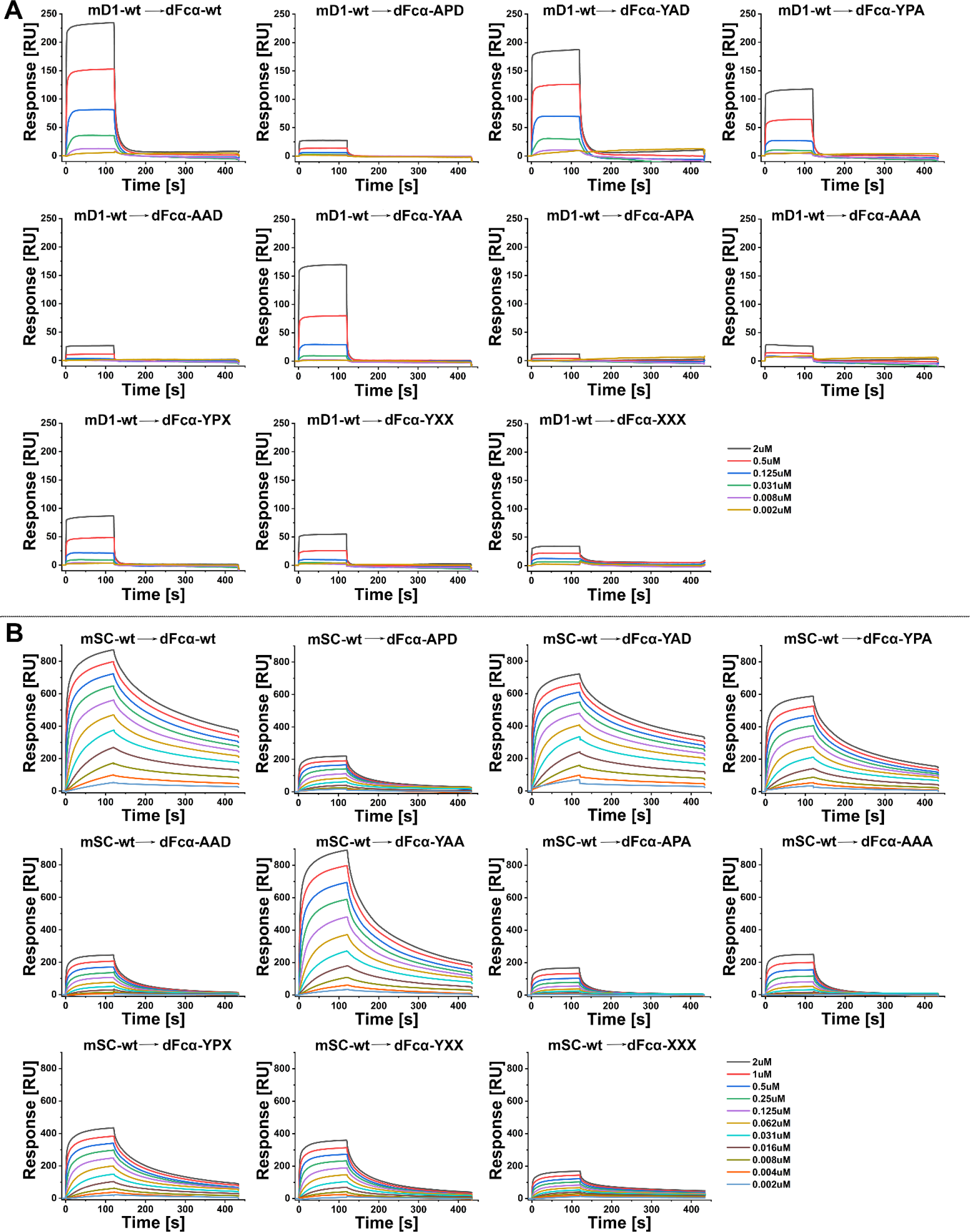
(A) SPR sensorgrams showing the responses for the entire dilution series of mD1-wt binding to the indicated dFcα variants (shown on the same scales). (B) SPR sensorgrams showing the responses for the entire dilution series of mSC-wt binding to the indicated dFcα variants (shown on the same scales). The responses for each analyte dilution are colored according to the keys (right). Results were consistent between 3 or more replicate experiments.

**Supplemental Figure-5:**
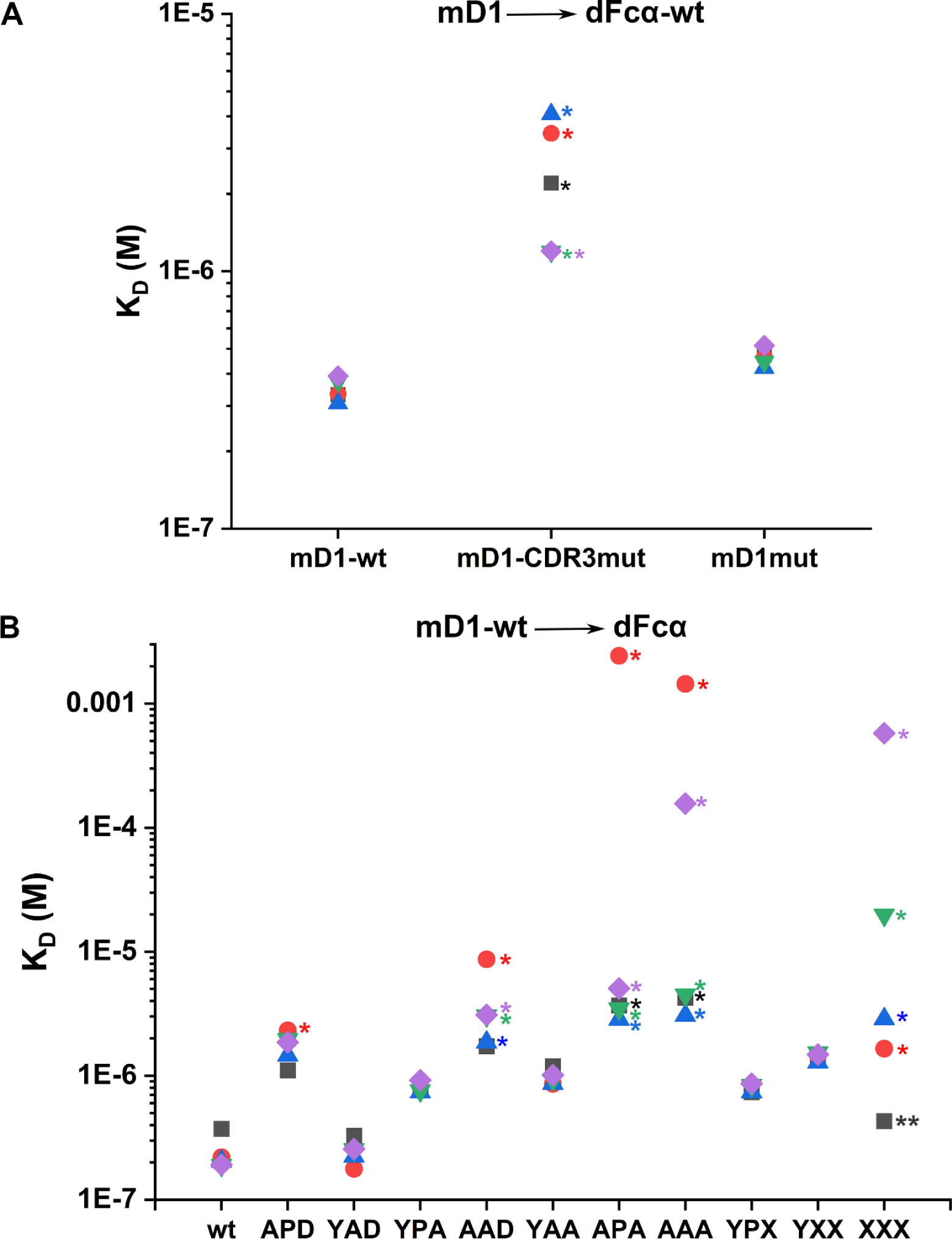
The steady state affinity K_D_ (M) values calculated from replicate experiments. (A) Reports the K_D_ values calculated from 5 replicate datasets (each represented by a unique colored shape) for mD1-wt, mD1-CDR3mut, and mD1-mut binding to dFcα-wt. The replicate data sets are from 5 different immobilized dFcα-wt spots titrated against a concentration gradient of mD1-wt, mD1-CDR3mut, mD1-mut. The green downward triangles represent the data in the main text Fig.2 and Fig.3. (B) Reports the K_D_ values calculated from 3-5 replicate experiments (each represented by a unique colored shape) for mD1-wt binding to dFcα-wt and mutants dFcα-APD, dFcα-YAD, dFcα-YPA, dFcα-AAD, dFcα-YAA, dFcα-APA, dFcα-AAA, dFcα-YPX, dFcα-YXX, dFcα-XXX. The black squares represent the data shown in the main text Fig.4. In both panels, * represents the data points for which the t-value (K_D_/ standard error) of steady state fitting was <10, and/or the K_D_ value was greater than 2μM (maximum analyte concentration), indicative of poor fit and low confidence in the calculated value. The black ** represents a K_D_ value calculated from steady state analysis with t=11 and low response for max concentration, hence low confidence in the calculated value, consistent with replicate experiments in which t<10.

## Supplementary Sequence File

This file contains the amino acid and DNA sequences for the proteins and complexes used in this study. Sequences include the signal peptides, which are naturally cleaved prior to secretion. The mSC variants all contain the C470A and C504A mutations (in addition to noted, introduced mutations). Sequences are named according to the main text.

### >mSC-wt

MRLYLFTLLVTVFSGVSTKSPIFGPQEVSSIEGDSVSITCYYPDTSVNRHTRKYWCRQGASGMCTTLISSNGYLSKEYSGRANLINFPENNTFVINIEQLTQDDTGSYKCGLGTSNRGLSFDVSLEVSQVPELPSDTHVYTKDIGRNVTIECPFKRENAPSKKSLCKKTNQSCELVIDSTEKVNPSYIGRAKLFMKGTDLTVFYVNISHLTHNDAGLYICQAGEGPSADKKNVDLQVLAPEPELLYKDLRSSVTFECDLGREVANEAKYLCRMNKETCDVIINTLGKRDPDFEGRILITPKDDNGRFSVLITGLRKEDAGHYQCGAHSSGLPQEGWPIQTWQLFVNEESTIPNRRSVVKGVTGGSVAIACPYNPKESSSLKYWCRWEGDGNGHCPVLVGTQAQVQEEYEGRLALFDQPGNGTYTVILNQLTTEDAGFYWCLTNGDSRWRTTIELQVAEATREPNLEVTPQNATAVLGETFTVSCHYPAKFYSQEKYWCKWSNKGCHILPSHDEGARQSSVSADQSSQLVSMTLNPVSKEDEGWYWCGVKQGQTYGETTAIYIAVEERGSHHHHHH*

### >mSC-wt

ATGAGGCTCTACTTGTTCACGCTCTTGGTAACTGTCTTTTCAGGGGTCTCCACAAAAAGCCCCATATTTGGTCCCCAGGAGGTGAGTAGTATAGAAGGCGACTCTGTTTCCATCACGTGCTACTACCCAGACACCTCTGTCAACCGGCACACCCGGAAATACTGGTGCCGACAAGGAGCCAGCGGCATGTGCACAACGCTCATCTCTTCAAATGGCTACCTCTCCAAGGAGTATTCAGGCAGAGCCAACCTCATCAACTTCCCAGAGAACAACACATTTGTGATTAACATTGAGCAGCTCACCCAGGACGACACTGGGAGCTACAAGTGTGGCCTGGGTACCAGTAACCGAGGCCTGTCCTTCGATGTCAGCCTGGAGGTCAGCCAGGTTCCTGAGTTGCCGAGTGACACCCACGTCTACACAAAGGACATAGGCAGAAATGTGACCATTGAATGCCCTTTCAAAAGGGAGAATGCTCCCAGCAAGAAATCCCTGTGTAAGAAGACAAACCAGTCCTGCGAACTTGTCATTGACTCTACTGAGAAGGTGAACCCCAGCTATATAGGCAGAGCAAAACTTTTTATGAAAGGGACCGACCTAACTGTATTCTATGTCAACATTAGTCACCTAACGCACAATGATGCTGGGCTGTACATCTGCCAAGCTGGAGAAGGTCCTAGTGCTGATAAGAAGAATGTTGACCTCCAGGTGCTAGCGCCTGAGCCAGAGCTGCTTTATAAAGACCTGAGGTCCTCAGTGACTTTTGAATGTGACCTGGGCCGTGAGGTGGCAAACGAGGCCAAATATCTGTGCCGGATGAATAAGGAAACCTGTGATGTGATCATTAACACCCTGGGGAAGAGGGATCCAGACTTTGAGGGCAGGATCCTGATAACCCCCAAGGATGACAATGGCCGCTTCAGTGTGTTGATCACAGGCCTGAGGAAGGAGGATGCAGGGCACTACCAGTGTGGAGCCCACAGTTCTGGTTTGCCTCAAGAAGGCTGGCCCATCCAGACTTGGCAACTCTTTGTCAATGAAGAGTCTACCATTCCCAATCGTCGCTCTGTTGTGAAGGGAGTCACAGGAGGCTCTGTGGCCATCGCCTGTCCCTATAACCCCAAGGAAAGCAGCAGCCTCAAGTACTGGTGTCGCTGGGAAGGGGACGGAAATGGACATTGCCCCGTGCTTGTGGGGACCCAGGCCCAGGTGCAAGAAGAGTATGAAGGCCGACTGGCACTGTTTGATCAGCCAGGCAATGGTACTTACACTGTCATCCTCAACCAGCTCACCACCGAGGATGCTGGCTTCTATTGGTGTCTTACCAATGGTGACTCTCGCTGGAGAACCACAATAGAACTCCAGGTTGCCGAAGCTACAAGGGAGCCAAACCTTGAGGTGACGCCACAGAACGCAACAGCAGTACTAGGAGAGACCTTCACCGTTTCCTGCCACTATCCGGCTAAATTCTACTCCCAGGAGAAATACTGGTGCAAGTGGAGCAACAAGGGTTGCCACATCCTGCCAAGCCATGACGAAGGTGCCCGCCAATCTTCTGTGAGCGCAGACCAGAGCAGCCAGCTGGTCTCCATGACCCTGAACCCGGTCAGTAAGGAAGATGAAGGCTGGTACTGGTGTGGGGTAAAGCAAGGCCAGACCTATGGAGAAACTACCGCCATCTATATAGCAGTTGAAGAGAGGgggtccCACCATCACCATCACCATTGATAG

### >mSC-D1-CDR3mut

MRLYLFTLLVTVFSGVSTKSPIFGPQEVSSIEGDSVSITCYYPDTSVNRHTRKYWCRQGASGMCTTLISSNGYLSKEYSGRANLINFPENNTFVINIEQLTQDDTGSYKCGLGAGGAGASFDVSLEVSQVPELPSDTHVYTKDIGRNVTIECPFKRENAPSKKSLCKKTNQSCELVIDSTEKVNPSYIGRAKLFMKGTDLTVFYVNISHL THNDAGLYICQAGEGPSADKKNVDLQVLAPEPELLYKDLRSSVTFECDLGREVANEAKYLCRMNKETCDVIINTLGKRDPDFEGRILITPKDDNGRFSVLITGLRKEDAGHYQCGAHSSGLPQEGWPIQTWQLFVNEESTIPNRRSVVKGVTGGSVAIACPYNPKESSSLKYWCRWEGDGNGHCPVLVGTQAQVQEEYEGRLALFDQPGNGTYTVILNQLTTEDAGFYWCLTNGDSRWRTTIELQVAEATREPNLEVTPQNATAVLGETFTVSCHYPAKFYSQEKYWCKWSNKGCHILPSHDEGARQSSVSADQSSQLVSMTLNPVSKEDEGWYWCGVKQGQTYGETTAIYIAVEERGSHHHHHH*

### >mSC-D1-CDR3mut

ATGAGGCTCTACTTGTTCACGCTCTTGGTAACTGTCTTTTCAGGGGTCTCCACAAAAAGCCCCATATTTGGTCCCCAGGAGGTGAGTAGTATAGAAGGCGACTCTGTTTCCATCACGTGCTACTACCCAGACACCTCTGTCAACCGGCACACCCGGAAATACTGGTGCCGACAAGGAGCCAGCGGCATGTGCACAACGCTCATCTCTTCAAATGGCTACCTCTCCAAGGAGTATTCAGGCAGAGCCAACCTCATCAACTTCCCAGAGAACAACACATTTGTGATTAACATTGAGCAGCTCACCCAGGACGACACTGGGAGCTACAAGTGTGGCCTGGGTGCAGGTGGCGCGGGCGCATCCTTCGATGTCAGCCTGGAGGTCAGCCAGGTTCCTGAGTTGCCGAGTGACACCCACGTCTACACAAAGGACATAGGCAGAAATGTGACCATTGAATGCCCTTTCAAAAGGGAGAATGCTCCCAGCAAGAAATCCCTGTGTAAGAAGACAAACCAGTCCTGCGAACTTGTCATTGACTCTACTGAGAAGGTGAACCCCAGCTATATAGGCAGAGCAAAACTTTTTATGAAAGGGACCGACCTAACTGTATTCTATGTCAACATTAGTCACCTAACGCACAATGATGCTGGGCTGTACATCTGCCAAGCTGGAGAAGGTCCTAGTGCTGATAAGAAGAATGTTGACCTCCAGGTGCTAGCGCCTGAGCCAGAGCTGCTTTATAAAGACCTGAGGTCCTCAGTGACTTTTGAATGTGACCTGGGCCGTGAGGTGGCAAACGAGGCCAAATATCTGTGCCGGATGAATAAGGAAACCTGTGATGTGATCATTAACACCCTGGGGAAGAGGGATCCAGACTTTGAGGGCAGGATCCTGATAACCCCCAAGGATGACAATGGCCGCTTCAGTGTGTTGATCACAGGCCTGAGGAAGGAGGATGCAGGGCACTACCAGTGTGGAGCCCACAGTTCTGGTTTGCCTCAAGAAGGCTGGCCCATCCAGACTTGGCAACTCTTTGTCAATGAAGAGTCTACCATTCCCAATCGTCGCTCTGTTGTGAAGGGAGTCACAGGAGGCTCTGTGGCCATCGCCTGTCCCTATAACCCCAAGGAAAGCAGCAGCCTCAAGTACTGGTGTCGCTGGGAAGGGGACGGAAATGGACATTGCCCCGTGCTTGTGGGGACCCAGGCCCAGGTGCAAGAAGAGTATGAAGGCCGACTGGCACTGTTTGATCAGCCAGGCAATGGTACTTACACTGTCATCCTCAACCAGCTCACCACCGAGGATGCTGGCTTCTATTGGTGTCTTACCAATGGTGACTCTCGCTGGAGAACCACAATAGAACTCCAGGTTGCCGAAGCTACAAGGGAGCCAAACCTTGAGGTGACGCCACAGAACGCAACAGCAGTACTAGGAGAGACCTTCACCGTTTCCTGCCACTATCCGGCTAAATTCTACTCCCAGGAGAAATACTGGTGCAAGTGGAGCAACAAGGGTTGCCACATCCTGCCAAGCCATGACGAAGGTGCCCGCCAATCTTCTGTGAGCGCAGACCAGAGCAGCCAGCTGGTCTCCATGACCCTGAACCCGGTCAGTAAGGAAGATGAAGGCTGGTACTGGTGTGGGGTAAAGCAAGGCCAGACCTATGGAGAAACTACCGCCATCTATATAGCAGTTGAAGAGAGGgggtccCACCATCACCATCACCATTGATAG

### >mSC-D1mut

MRLYLFTLLVTVFSGVSTKSPIAGPQEVSSIEGDSVSITCAYADTSVNRHTRKYWCRQGASGMCTTLISSNGYLSKEYSGRANLINFPENNTFVINIEQLTQDDTGSYKCGLGTSNRGLSFDVSLEVSQVPELPSDTHVYTKDIGRNVTIECPFKRENAPSKKSLCKKTNQSCELVIDSTEKVNPSYIGRAKLFMKGTDLTVFYVNISHLTHNDAGLYICQAGEGPSADKKNVDLQVLAPEPELLYKDLRSSVTFECDLGREVANEAKYLCRMNKETCDVIINTLGKRDPDFEGRILITPKDDNGRFSVLITGLRKEDAGHYQCGAHSSGLPQEGWPIQTWQLFVNEESTIPNRRSVVKGVTGGSVAIACPYNPKESSSLKYWCRWEGDGNGHCPVLVGTQAQVQEEYEGRLALFDQPGNGTYTVILNQLTTEDAGFYWCLTNGDSRWRTTIELQVAEATREPNLEVTPQNATAVLGETFTVSCHYPAKFYSQEKYWCKWSNKGCHILPSHDEGARQSSVSADQSSQLVSMTLNPVSKEDEGWYWCGVKQGQTYGETTAIYIAVEERGSHHHHHH*

### >mSC-D1mut

ATGAGGCTCTACTTGTTCACGCTCTTGGTAACTGTCTTTTCAGGGGTCTCCACAAAAAGCCCCATAGCAGGTCCCCAGGAGGTGAGTAGTATAGAAGGCGACTCTGTTTCCATCACGTGCGCATACGCAGACACCTCTGTCAACCGGCACACCCGGAAATACTGGTGCCGACAAGGAGCCAGCGGCATGTGCACAACGCTCATCTCTTCAAATGGCTACCTCTCCAAGGAGTATTCAGGCAGAGCCAACCTCATCAACTTCCCAGAGAACAACACATTTGTGATTAACATTGAGCAGCTCACCCAGGACGACACTGGGAGCTACAAGTGTGGCCTGGGTACCAGTAACCGAGGCCTGTCCTTCGATGTCAGCCTGGAGGTCAGCCAGGTTCCTGAGTTGCCGAGTGACACCCACGTCTACACAAAGGACATAGGCAGAAATGTGACCATTGAATGCCCTTTCAAAAGGGAGAATGCTCCCAGCAAGAAATCCCTGTGTAAGAAGACAAACCAGTCCTGCGAACTTGTCATTGACTCTACTGAGAAGGTGAACCCCAGCTATATAGGCAGAGCAAAACTTTTTATGAAAGGGACCGACCTAACTGTATTCTATGTCAACATTAGTCACCTAACGCACAATGATGCTGGGCTGTACATCTGCCAAGCTGGAGAAGGTCCTAGTGCTGATAAGAAGAATGTTGACCTCCAGGTGCTAGCGCCTGAGCCAGAGCTGCTTTATAAAGACCTGAGGTCCTCAGTGACTTTTGAATGTGACCTGGGCCGTGAGGTGGCAAACGAGGCCAAATATCTGTGCCGGATGAATAAGGAAACCTGTGATGTGATCATTAACACCCTGGGGAAGAGGGATCCAGACTTTGAGGGCAGGATCCTGATAACCCCCAAGGATGACAATGGCCGCTTCAGTGTGTTGATCACAGGCCTGAGGAAGGAGGATGCAGGGCACTACCAGTGTGGAGCCCACAGTTCTGGTTTGCCTCAAGAAGGCTGGCCCATCCAGACTTGGCAACTCTTTGTCAATGAAGAGTCTACCATTCCCAATCGTCGCTCTGTTGTGAAGGGAGTCACAGGAGGCTCTGTGGCCATCGCCTGTCCCTATAACCCCAAGGAAAGCAGCAGCCTCAAGTACTGGTGTCGCTGGGAAGGGGACGGAAATGGACATTGCCCCGTGCTTGTGGGGACCCAGGCCCAGGTGCAAGAAGAGTATGAAGGCCGACTGGCACTGTTTGATCAGCCAGGCAATGGTACTTACACTGTCATCCTCAACCAGCTCACCACCGAGGATGCTGGCTTCTATTGGTGTCTTACCAATGGTGACTCTCGCTGGAGAACCACAATAGAACTCCAGGTTGCCGAAGCTACAAGGGAGCCAAACCTTGAGGTGACGCCACAGAACGCAACAGCAGTACTAGGAGAGACCTTCACCGTTTCCTGCCACTATCCGGCTAAATTCTACTCCCAGGAGAAATACTGGTGCAAGTGGAGCAACAAGGGTTGCCACATCCTGCCAAGCCATGACGAAGGTGCCCGCCAATCTTCTGTGAGCGCAGACCAGAGCAGCCAGCTGGTCTCCATGACCCTGAACCCGGTCAGTAAGGAAGATGAAGGCTGGTACTGGTGTGGGGTAAAGCAAGGCCAGACCTATGGAGAAACTACCGCCATCTATATAGCAGTTGAAGAGAGGgggtccCACCATCACCATCACCATTGATAG

### >mSC-D3mut

MRLYLFTLLVTVFSGVSTKSPIFGPQEVSSIEGDSVSITCYYPDTSVNRHTRKYWCRQGASGMCTTLISSNGYLSKEYSGRANLINFPENNTFVINIEQLTQDDTGSYKCGLGTSNRGLSFDVSLEVSQVPELPSDTHVYTKDIGRNVTIECPFKRENAPSKKSLCKKTNQSCELVIDSTEKVNPSYIGRAKLFMKGTDLTVFYVNISHLTHNDAGLYICQAGEGPSADKKNVDLQVLAPEPEALAKDLRSSVTFECDLGREVANEAKYLCRMNKETCDVIINTLGKRDPDFEGRILITPKDDNGRFSVLITGLRKEDAGHYQCGAHSSGLPQEGWPIQTWQLAVNEESTIPNRRSVVKGVTGGSVAIACPYNPKESSSLKYWCRWEGDGNGHCPVLVGTQAQVQEEYEGRLALFDQPGNGTYTVILNQLTTEDAGFYWCLTNGDSRWRTTIELQVAEATREPNLEVTPQNATAVLGETFTVSCHYPAKFYSQEKYWCKWSNKGCHILPSHDEGARQSSVSADQSSQLVSMTLNPVSKEDEGWYWCGVKQGQTYGETTAIYIAVEERGSHHHHHH*

### >mSC-D3mut

ATGAGGCTCTACTTGTTCACGCTCTTGGTAACTGTCTTTTCAGGGGTCTCCACAAAAAGCCCCATATTTGGTCCCCAGGAGGTGAGTAGTATAGAAGGCGACTCTGTTTCCATCACGTGCTACTACCCAGACACCTCTGTCAACCGGCACACCCGGAAATACTGGTGCCGACAAGGAGCCAGCGGCATGTGCACAACGCTCATCTCTTCAAATGGCTACCTCTCCAAGGAGTATTCAGGCAGAGCCAACCTCATCAACTTCCCAGAGAACAACACATTTGTGATTAACATTGAGCAGCTCACCCAGGACGACACTGGGAGCTACAAGTGTGGCCTGGGTACCAGTAACCG AGGCCTGTCCTTCGATGTCAGCCTGGAGGTCAGCCAGGTTCCTGAGTTGCCGAGTGACACCCACGTCTACACAAAGGACATAGGCAGAAATGTGACCATTGAATGCCCTTTCAAAAGGGAGAATGCTCCCAGCAAGAAATCCCTGTGTAAGAAGACAAACCAGTCCTGCGAACTTGTCATTGACTCTACTGAGAAGGTGAACCCCAGCTATATAGGCAGAGCAAAACTTTTTATGAAAGGGACCGACCTAACTGTATTCTATGTCAACATTAGTCACCTAACGCACAATGATGCTGGGCTGTACATCTGCCAAGCTGGAGAAGGTCCTAGTGCTGATAAGAAGAATGTTGACCTCCAGGTGCTAGCGCCTGAGCCAGAGGCACTTGCAAAAGACCTGAGGTCCTCAGTGACTTTTGAATGTGACCTGGGCCGTGAGGTGGCAAACGAGGCCAAATATCTGTGCCGGATGAATAAGGAAACCTGTGATGTGATCATTAACACCCTGGGGAAGAGGGATCCAGACTTTGAGGGCAGGATCCTGATAACCCCCAAGGATGACAATGGCCGCTTCAGTGTGTTGATCACAGGCCTGAGGAAGGAGGATGCAGGGCACTACCAGTGTGGAGCCCACAGTTCTGGTTTGCCTCAAGAAGGCTGGCCCATCCAGACTTGGCAACTCGCAGTCAATGAAGAGTCTACCATTCCCAATCGTCGCTCTGTTGTGAAGGGAGTCACAGGAGGCTCTGTGGCCATCGCCTGTCCCTATAACCCCAAGGAAAGCAGCAGCCTCAAGTACTGGTGTCGCTGGGAAGGGGACGGAAATGGACATTGCCCCGTGCTTGTGGGGACCCAGGCCCAGGTGCAAGAAGAGTATGAAGGCCGACTGGCACTGTTTGATCAGCCAGGCAATGGTACTTACACTGTCATCCTCAACCAGCTCACCACCGAGGATGCTGGCTTCTATTGGTGTCTTACCAATGGTGACTCTCGCTGGAGAACCACAATAGAACTCCAGGTTGCCGAAGCTACAAGGGAGCCAAACCTTGAGGTGACGCCACAGAACGCAACAGCAGTACTAGGAGAGACCTTCACCGTTTCCTGCCACTATCCGGCTAAATTCTACTCCCAGGAGAAATACTGGTGCAAGTGGAGCAACAAGGGTTGCCACATCCTGCCAAGCCATGACGAAGGTGCCCGCCAATCTTCTGTGAGCGCAGACCAGAGCAGCCAGCTGGTCTCCATGACCCTGAACCCGGTCAGTAAGGAAGATGAAGGCTGGTACTGGTGTGGGGTAAAGCAAGGCCAGACCTATGGAGAAACTACCGCCATCTATATAGCAGTTGAAGAGAGGgggtccCACCATCACCATCACCATTGATAG

### >mSC-D1-D3mut

MRLYLFTLLVTVFSGVSTKSPIAGPQEVSSIEGDSVSITCAYADTSVNRHTRKYWCRQGASGMCTTLISSNGYLSKEYSGRANLINFPENNTFVINIEQLTQDDTGSYKCGLGTSNRGLSFDVSLEVSQVPELPSDTHVYTKDIGRNVTIECPFKRENAPSKKSLCKKTNQSCELVIDSTEKVNPSYIGRAKLFMKGTDLTVFYVNISHLTHNDAGLYICQAGEGPSADKKNVDLQVLAPEPEALAKDLRSSVTFECDLGREVANEAKYLCRMNKETCDVIINTLGKRDPDFEGRILITPKDDNGRFSVLITGLRKEDAGHYQCGAHSSGLPQEGWPIQTWQLAVNEESTIPNRRSVVKGVTGGSVAIACPYNPKESSSLKYWCRWEGDGNGHCPVLVGTQAQVQEEYEGRLALFDQPGNGTYTVILNQLTTEDAGFYWCLTNGDSRWRTTIELQVAEATREPNLEVTPQNATAVLGETFTVSCHYPAKFYSQEKYWCKWSNKGCHILPSHDEGARQSSVSADQSSQLVSMTLNPVSKEDEGWYWCGVKQGQTYGETTAIYIAVEERGSHHHHHH*

### >mSC-D1-D3mut

ATGAGGCTCTACTTGTTCACGCTCTTGGTAACTGTCTTTTCAGGGGTCTCCACAAAAAGCCCCATAGCAGGTCCCCAGGAGGTGAGTAGTATAGAAGGCGACTCTGTTTCCATCACGTGCGCATACGCAGACACCTCTGTCAACCGGCACACCCGGAAATACTGGTGCCGACAAGGAGCCAGCGGCATGTGCACAACGCTCATCTCTTCAAATGGCTACCTCTCCAAGGAGTATTCAGGCAGAGCCAACCTCATCAACTTCCCAGAGAACAACACATTTGTGATTAACATTGAGCAGCTCACCCAGGACGACACTGGGAGCTACAAGTGTGGCCTGGGTACCAGTAACCGAGGCCTGTCCTTCGATGTCAGCCTGGAGGTCAGCCAGGTTCCTGAGTTGCCGAGTGACACCCACGTCTACACAAAGGACATAGGCAGAAATGTGACCATTGAATGCCCTTTCAAAAGGGAGAATGCTCCCAGCAAGAAATCCCTGTGTAAGAAGACAAACCAGTCCTGCGAACTTGTCATTGACTCTACTGAGAAGGTGAACCCCAGCTATATAGGCAGAGCAAAACTTTTTATGAAAGGGACCGACCTAACTGTATTCTATGTCAACATTAGTCACCTAACGCACAATGATGCTGGGCTGTACATCTGCCAAGCTGGAGAAGGTCCTAGTGCTGATAAGAAGAATGTTGACCTCCAGGTGCTAGCGCCTGAGCCAGAGGCACTTGCAAAAGACCTGAGGTCCTCAGTGACTTTTGAATG TGACCTGGGCCGTGAGGTGGCAAACGAGGCCAAATATCTGTGCCGGATGAATAAGGAAACCTGTGATGTGATCATTAACACCCTGGGGAAGAGGGATCCAGACTTTGAGGGCAGGATCCTGATAACCCCCAAGGATGACAATGGCCGCTTCAGTGTGTTGATCACAGGCCTGAGGAAGGAGGATGCAGGGCACTACCAGTGTGGAGCCCACAGTTCTGGTTTGCCTCAAGAAGGCTGGCCCATCCAGACTTGGCAACTCGCAGTCAATGAAGAGTCTACCATTCCCAATCGTCGCTCTGTTGTGAAGGGAGTCACAGGAGGCTCTGTGGCCATCGCCTGTCCCTATAACCCCAAGGAAAGCAGCAGCCTCAAGTACTGGTGTCGCTGGGAAGGGGACGGAAATGGACATTGCCCCGTGCTTGTGGGGACCCAGGCCCAGGTGCAAGAAGAGTATGAAGGCCGACTGGCACTGTTTGATCAGCCAGGCAATGGTACTTACACTGTCATCCTCAACCAGCTCACCACCGAGGATGCTGGCTTCTATTGGTGTCTTACCAATGGTGACTCTCGCTGGAGAACCACAATAGAACTCCAGGTTGCCGAAGCTACAAGGGAGCCAAACCTTGAGGTGACGCCACAGAACGCAACAGCAGTACTAGGAGAGACCTTCACCGTTTCCTGCCACTATCCGGCTAAATTCTACTCCCAGGAGAAATACTGGTGCAAGTGGAGCAACAAGGGTTGCCACATCCTGCCAAGCCATGACGAAGGTGCCCGCCAATCTTCTGTGAGCGCAGACCAGAGCAGCCAGCTGGTCTCCATGACCCTGAACCCGGTCAGTAAGGAAGATGAAGGCTGGTACTGGTGTGGGGTAAAGCAAGGCCAGACCTATGGAGAAACTACCGCCATCTATATAGCAGTTGAAGAGAGGgggtccCACCATCACCATCACCATTGATAG

### >mD1-wt

MRLYLFTLLVTVFSGVSTKSPIFGPQEVSSIEGDSVSITCYYPDTSVNRHTRKYWCRQGASGMCTTLISSNGYLSKEYSGRANLINFPENNTFVINIEQLTQDDTGSYKCGLGTSNRGLSFDVSLEVSQVPELPSDTHVYTKDIGRNGSHHHHHH*

### >mD1-wt

ATGAGGCTCTACTTGTTCACGCTCTTGGTAACTGTCTTTTCAGGGGTCTCCACAAAAAGCCCCATATTTGGTCCCCAGGAGGTGAGTAGTATAGAAGGCGACTCTGTTTCCATCACGTGCTACTACCCAGACACCTCTGTCAACCGGCACACCCGGAAATACTGGTGCCGACAAGGAGCCAGCGGCATGTGCACAACGCTCATCTCTTCAAATGGCTACCTCTCCAAGGAGTATTCAGGCAGAGCCAACCTCATCAACTTCCCAGAGAACAACACATTTGTGATTAACATTGAGCAGCTCACCCAGGACGACACTGGGAGCTACAAGTGTGGCCTGGGTACCAGTAACCGAGGCCTGTCCTTCGATGTCAGCCTGGAGGTCAGCCAGGTTCCTGAGTTGCCGAGTGACACCCACGTCTACACAAAGGACATAGGCAGAAATgggtccCACCATCACCATCACCATTGA

### >mD1-CDR3mut

MRLYLFTLLVTVFSGVSTKSPIFGPQEVSSIEGDSVSITCYYPDTSVNRHTRKYWCRQGASGMCTTLISSNGYLSKEYSGRANLINFPENNTFVINIEQLTQDDTGSYKCGLGAGGAGASFDVSLEVSQVPELPSDTHVYTKDIGRNGSHHHHHH*

### >mD1-CDR3mut

ATGAGGCTCTACTTGTTCACGCTCTTGGTAACTGTCTTTTCAGGGGTCTCCACAAAAAGCCCCATATTTGGTCCCCAGGAGGTGAGTAGTATAGAAGGCGACTCTGTTTCCATCACGTGCTACTACCCAGACACCTCTGTCAACCGGCACACCCGGAAATACTGGTGCCGACAAGGAGCCAGCGGCATGTGCACAACGCTCATCTCTTCAAATGGCTACCTCTCCAAGGAGTATTCAGGCAGAGCCAACCTCATCAACTTCCCAGAGAACAACACATTTGTGATTAACATTGAGCAGCTCACCCAGGACGACACTGGGAGCTACAAGTGTGGCCTGGGTGCAGGTGGCGC GGGCGCATCCTTCGATGTCAGCCTGGAGGTCAGCCAGGTTCCTGAGTTGCCGAGTGACACCCACGTCTACACAAAGGACATAGGCAGAAATgggtccCACCATCACCATCACCATTGA

### >mD1-mut

MRLYLFTLLVTVFSGVSTKSPIAGPQEVSSIEGDSVSITCAYADTSVNRHTRKYWCRQGASGMCTTLISSNGYLSKEYSGRANLINFPENNTFVINIEQLTQDDTGSYKCGLGTSNRGLSFDVSLEVSQVPELPSDTHVYTKDIGRNGSHHHHHH*

### >mD1-mut

ATGAGGCTCTACTTGTTCACGCTCTTGGTAACTGTCTTTTCAGGGGTCTCCACAAAAAGCCCCATAGCAGGTCCCCAGGAGGTGAGTAGTATAGAAGGCGACTCTGTTTCCATCACGTGCGCATACGCAGACACCTCTGTCAACCGGCACACCCGGAAATACTGGTGCCGACAAGGAGCCAGCGGCATGTGCACAACGCTCATCTCTTCAAATGGCTACCTCTCCAAGGAGTATTCAGGCAGAGCCAACCTCATCAACTTCCCAGAGAACAACACATTTGTGATTAACATTGAGCAGCTCACCCAGGACGACACTGGGAGCTACAAGTGTGGCCTGGGTACCAGTAACCGAGGCCTGTCCTTCGATGTCAGCCTGGAGGTCAGCCAGGTTCCTGAGTTGCCGAGTGACACCCACGTCTACACAAAGGACATAGGCAGAAATgggtccCACCATCACCATCACCATTGA

### >mJC-wt

MKTHLLLWGVLAIFVKAVLVTGDDEATILADNKCMCTRVTSRIIPSTEDPNEDIVERNIRIVVPLNNRENISDPTSPLRRNFVYHLSDVCKKCDPVEVELEDQVVTATQSNICNEDDGVPETCYMYDRNKCYTTMVPLRYHGETKMVQAALTPDSCYPD*

### >mJC-wt

ATGAAGACCCACCTGCTTCTCTGGGGAGTCCTGGCCATTTTTGTTAAGGCTGTCCTTGTAACAGGTGACGACGAAGCGACCATTCTTGCTGACAACAAATGCATGTGTACCCGAGTTACCTCTAGGATCATCCCTTCCACCGAGGATCCTAATGAGGACATTGTGGAGAGAAATATCCGAATTGTTGTCCCTTTGAACAACAGGGAGAATATCTCTGATCCCACCTCCCCACTGAGAAGGAACTTTGTATACCATTTGTCAGACGTCTGTAAGAAATGCGATCCTGTGGAAGTGGAGCTGGAAGATCAGGTTGTTACTGCCACCCAGAGCAACATCTGCAATGAAGACGATGGTGTTCCTGAGACCTGCTACATGTATGACAGAAACAAGTGCTATACCACTATGGTCCCACTTAGGTATCATGGTGAGACCAAAATGGTGCAAGCAGCCTTGACCCCCGATTCTTGCTACCCTGACTAGTAA

### >mJC-YPA

MKTHLLLWGVLAIFVKAVLVTGDDEATILADNKCMCTRVTSRIIPSTEDPNEDIVERNIRIVVPLNNRENISDPTSPLRRNFVYHLSDVCKKCDPVEVELEDQVVTATQSNICNEDDGVPETCYMYDRNKCYTTMVPLRYHGETKMVQAALTPDSCYPA*

### >mJC-YPA

ATGAAGACCCACCTGCTTCTCTGGGGAGTCCTGGCCATTTTTGTTAAGGCTGTCCTTGTAACAGGTGACGACGAAGCGACCATTCTTGCTGACAACAAATGCATGTGTACCCGAGTTACCTCTAGGATCATCCCTTCCACCGAGGATCCTAATGAGGACATTGTGGAGAGAAATATCCGAATTGTTGTCCCTTTGAACAACAGGGAGAATATCTCTGATCCCACCTCCCCACTGAGAAGGAACTTTGTATACCATTTGTCAGACGTCTGTAAGAAATGCGATCCTGTGGAAGTGGAGCTGGAAGATCAGGTTGTTACTGCCACCCAGAGCAACATCTGCAATGAAGACGATGGTGTTCCTGAGACCTGCTACATGTATGACAGAAACAAGTGCTATACCACTATGGTCCCACTTAGGTATCATGGTGAGACCAAAATGGTGCAAGCAGCCTTGACCCCCGATTCTTGCTACCCTGCCTAG

### >mJC-YAD

MKTHLLLWGVLAIFVKAVLVTGDDEATILADNKCMCTRVTSRIIPSTEDPNEDIVERNIRIVVPLNNRENISDPTSPLRRNFVYHLSDVCKKCDPVEVELEDQVVTATQSNICNEDDGVPETCYMYDRNKCYTTMVPLRYHGETKMVQAALTPDSCYAD*

### >mJC-YAD

ATGAAGACCCACCTGCTTCTCTGGGGAGTCCTGGCCATTTTTGTTAAGGCTGTCCTTGTAACAGGTGACGACGAAGCGACCATTCTTGCTGACAACAAATGCATGTGTACCCGAGTTACCTCTAGGATCATCCCTTCCACCGAGGATCCTAATGAGGACATTGTGGAGAGAAATATCCGAATTGTTGTCCCTTTGAACAACAGGGAGAATATCTCTGATCCCACCTCCCCACTGAGAAGGAACTTTGTATACCATTTGTCAGACGTCTGTAAGAAATGCGATCCTGTGGAAGTGGAGCTGGAAGATCAGGTTGTTACTGCCACCCAGAGCAACATCTGCAATGAAGACGATGGTGTTCCTGAGACCTGCTACATGTATGACAGAAACAAGTGCTATACCACTATGGTCCCACTTAGGTATCATGGTGAGACCAAAATGGTGCAAGCAGCCTTGACCCCCGATTCTTGCTACGCCGACTAG

### >mJC-APD

MKTHLLLWGVLAIFVKAVLVTGDDEATILADNKCMCTRVTSRIIPSTEDPNEDIVERNIRIVVPLNNRENISDPTSPLRRNFVYHLSDVCKKCDPVEVELEDQVVTATQSNICNEDDGVPETCYMYDRNKCYTTMVPLRYHGETKMVQAALTPDSCAPD*

### >mJC-APD

ATGAAGACCCACCTGCTTCTCTGGGGAGTCCTGGCCATTTTTGTTAAGGCTGTCCTTGTAACAGGTGACGACGAAGCGACCATTCTTGCTGACAACAAATGCATGTGTACCCGAGTTACCTCTAGGATCATCCCTTCCACCGAGGATCCTAATGAGGACATTGTGGAGAGAAATATCCGAATTGTTGTCCCTTTGAACAACAGGGAGAATATCTCTGATCCCACCTCCCCACTGAGAAGGAACTTTGTATACCATTTGTCAGACGTCTGTAAGAAATGCGATCCTGTGGAAGTGGAGCTGGAAGATCAGGTTGTTACTGCCACCCAGAGCAACATCTGCAATGAAGACGATGGTGTTCCTGAGACCTGCTACATGTATGACAGAAACAAGTGCTATACCACTATGGTCCCACTTAGGTATCATGGTGAGACCAAAATGGTGCAAGCAGCCTTGACCCCCGATTCTTGCGCCCCTGACTAG

### >mJC-YAA

MKTHLLLWGVLAIFVKAVLVTGDDEATILADNKCMCTRVTSRIIPSTEDPNEDIVERNIRIVVPLNNRENISDPTSPLRRNFVYHLSDVCKKCDPVEVELEDQVVTATQSNICNEDDGVPETCYMYDRNKCYTTMVPLRYHGETKMVQAALTPDSCYAA*

### >mJC-YAA

ATGAAGACCCACCTGCTTCTCTGGGGAGTCCTGGCCATTTTTGTTAAGGCTGTCCTTGTAACAGGTGACGACGAAGCGACCATTCTTGCTGACAACAAATGCATGTGTACCCGAGTTACCTCTAGGATCATCCCTTCCACCGAGGATCCTAATGAGGACATTGTGGAGAGAAATATCCGAATTGTTGTCCCTTTGAACAACAGGGAGAATATCTCTGATCCCACCTCCCCACTGAGAAGGAACTTTGTATACCATTTGTCAGACGTCTGTAAGAAATGCGATCCTGTGGAAGTGGAGCTGGAAGATCAGGTTGTTACTGCCACCCAGAGCAACATCTGCAATGAAGACGATGGTGTTCCTGAGACCTGCTACATGTATGACAGAAACAAGTGCTATACCACTATGGTCCCACTTAGGTATCATGGTGAGACCAAAATGGTGCAAGCAGCCTTGACCCCCGATTCTTGCTACGCTGCCTAG

### >mJC-APA

MKTHLLLWGVLAIFVKAVLVTGDDEATILADNKCMCTRVTSRIIPSTEDPNEDIVERNIRIVVPLNNRENISDPTSPLRRNFVYHLSDVCKKCDPVEVELEDQVVTATQSNICNEDDGVPETCYMYDRNKCYTTMVPLRYHGETKMVQAALTPDSCAPA*

### >mJC-APA

ATGAAGACCCACCTGCTTCTCTGGGGAGTCCTGGCCATTTTTGTTAAGGCTGTCCTTGTAACAGGTGACGACGAAGCGACCATTCTTGCTGACAACAAATGCATGTGTACCCGAGTTACCTCTAGGATCATCCCTTCCACCGAGGATCCTAATGAGGACATTGTGGAGAGAAATATCCGAATTGTTGTCCCTTTGAACAACAGGGAGAATATCTCTGATCCCACCTCCCCACTGAGAAGGAACTTTGTATACCATTTGTCAGACGTCTGTAAGAAATGCGATCCTGTGGAAGTGGAGCTGGAAGATCAGGTTGTTACTGCCACCCAGAGCAACATCTGCAATGAAGACGATGGTGTTCCTGAGACCTGCTACATGTATGACAGAAACAAGTGCTATACCACTATGGTCCCACTTAGGTATCATGGTGAGACCAAAATGGTGCAAGCAGCCTTGACCCCCGATTCTTGCGCCCCTGCCTAG

### >mJC-AAD

MKTHLLLWGVLAIFVKAVLVTGDDEATILADNKCMCTRVTSRIIPSTEDPNEDIVERNIRIVVPLNNRENISDPTSPLRRNFVYHLSDVCKKCDPVEVELEDQVVTATQSNICNEDDGVPETCYMYDRNKCYTTMVPLRYHGETKMVQAALTPDSCAAD*

### >mJC-AAD

ATGAAGACCCACCTGCTTCTCTGGGGAGTCCTGGCCATTTTTGTTAAGGCTGTCCTTGTAACAGGTGACGACGAAGCGACCATTCTTGCTGACAACAAATGCATGTGTACCCGAGTTACCTCTAGGATCATCCCTTCCACCGAGGATCCTAATGAGGACATTGTGGAGAGAAATATCCGAATTGTTGTCCCTTTGAACAACAGGGAGAATATCTCTGATCCCACCTCCCCACTGAGAAGGAACTTTGTATACCATTTGTCAGACGTCTGTAAGAAATGCG

ATCCTGTGGAAGTGGAGCTGGAAGATCAGGTTGTTACTGCCACCCAGAGCAACATCTGCAATGAAGACGATGGTGTTCCTGAGACCTGCTACATGTATGACAGAAACAAGTGCTATACCACTATGGTCCCACTTAGGTATCATGGTGAGACCAAAATGGTGCAAGCAGCCTTGACCCCCGATTCTTGCGCCGCTGACTAG

### >mJC-AAA

MKTHLLLWGVLAIFVKAVLVTGDDEATILADNKCMCTRVTSRIIPSTEDPNEDIVERNIRIVVPLNNRENISDPTSPLRRNFVYHLSDVCKKCDPVEVELEDQVVTATQSNICNEDDGVPETCYMYDRNKCYTTMVPLRYHGETKMVQAALTPDSCAAA*

### >mJC-AAA

ATGAAGACCCACCTGCTTCTCTGGGGAGTCCTGGCCATTTTTGTTAAGGCTGTCCTTGTAACAGGTGACGACGAAGCGACCATTCTTGCTGACAACAAATGCATGTGTACCCGAGTTACCTCTAGGATCATCCCTTCCACCGAGGATCCTAATGAGGACATTGTGGAGAGAAATATCCGAATTGTTGTCCCTTTGAACAACAGGGAGAATATCTCTGATCCCACCTCCCCACTGAGAAGGAACTTTGTATACCATTTGTCAGACGTCTGTAAGAAATGCGATCCTGTGGAAGTGGAGCTGGAAGATCAGGTTGTTACTGCCACCCAGAGCAACATCTGCAATGAAGACGATGGTGTTCCTGAGACCTGCTACATGTATGACAGAAACAAGTGCTATACCACTATGGTCCCACTTAGGTATCATGGTGAGACCAAAATGGTGCAAGCAGCCTTGACCCCCGATTCTTGCGCCGCTGCCTAG

### >mJC-YPX

MKTHLLLWGVLAIFVKAVLVTGDDEATILADNKCMCTRVTSRIIPSTEDPNEDIVERNIRIVVPLNNRENISDPTSPLRRNFVYHLSDVCKKCDPVEVELEDQVVTATQSNICNEDDGVPETCYMYDRNKCYTTMVPLRYHGETKMVQAALTPDSCYP*

### >mJC-YPX

ATGAAGACCCACCTGCTTCTCTGGGGAGTCCTGGCCATTTTTGTTAAGGCTGTCCTTGTAACAGGTGACGACGAAGCGACCATTCTTGCTGACAACAAATGCATGTGTACCCGAGTTACCTCTAGGATCATCCCTTCCACCGAGGATCCTAATGAGGACATTGTGGAGAGAAATATCCGAATTGTTGTCCCTTTGAACAACAGGGAGAATATCTCTGATCCCACCTCCCCACTGAGAAGGAACTTTGTATACCATTTGTCAGACGTCTGTAAGAAATGCGATCCTGTGGAAGTGGAGCTGGAAGATCAGGTTGTTACTGCCACCCAGAGCAACATCTGCAATGAAGACGATGGTGTTCCTGAGACCTGCTACATGTATGACAGAAACAAGTGCTATACCACTATGGTCCCACTTAGGTATCATGGTGAGACCAAAATGGTGCAAGCAGCCTTGACCCCCGATTCTTGCTACCCTTAG

### >mJC-YXX

MKTHLLLWGVLAIFVKAVLVTGDDEATILADNKCMCTRVTSRIIPSTEDPNEDIVERNIRIVVPLNNRENISDPTSPLRRNFVYHLSDVCKKCDPVEVELEDQVVTATQSNICNEDDGVPETCYMYDRNKCYTTMVPLRYHGETKMVQAALTPDSCY*

### >mJC-YXX

ATGAAGACCCACCTGCTTCTCTGGGGAGTCCTGGCCATTTTTGTTAAGGCTGTCCTTGTAACAGGTGACGACGAAGCGACCATTCTTGCTGACAACAAATGCATGTGTACCCGAGTTACCTCTAGGATCATCCCTTCCACCGAGGATCCTAATGAGGACATTGTGGAGAGAAATATCCGAATTGTTGTCCCTTTGAACAACAGGGAGAATATCTCTGATCCCACCTCCCCACTGAGAAGGAACTTTGTATACCATTTGTCAGACGTCTGTAAGAAATGCGATCCTGTGGAAGTGGAGCTGGAAGATCAGGTTGTTACTGCCACCCAGAGCAACATCTGCAATGAAGACGATGGTGTTCCTGAGACCTGCTACATGTATGACAGAAACAAGTGCTATACCACTATGGTCCCACTTAGGTATCATGGTGAGACCAAAATGGTGCAAGCAGCCTTGACCCCCGATTCTTGCTACTAG

### >mJC-XXX

MKTHLLLWGVLAIFVKAVLVTGDDEATILADNKCMCTRVTSRIIPSTEDPNEDIVERNIRIVVPLNNRENISDPTSPLRRNFVYHLSDVCKKCDPVEVELEDQVVTATQSNICNEDDGVPETCYMYDRNKCYTTMVPLRYHGETKMVQAALTPDSC*

### >mJC-XXX

ATGAAGACCCACCTGCTTCTCTGGGGAGTCCTGGCCATTTTTGTTAAGGCTGTCCTTGTAACAGGTGACGACGAAGCGACCATTCTTGCTGACAACAAATGCATGTGTACCCGAGTTACCTCTAGGATCATCCCTTCCACCGAGGATCCTAATGAGGACATTGTGGAGAGAAATATCCGAATTGTTGTCCCTTTGAACAACAGGGAGAATATCTCTGATCCCACCTCCCCACTGAGAAGGAACTTTGTATACCATTTGTCAGACGTCTGTAAGAAATGCGATCCTGTGGAAGTGGAGCTGGAAGATCAGGTTGTTACTGCCACCCAGAGCAACATCTGCAATGAAGACGATGGTGTTCCTGAGACCTGCTACATGTATGACAGAAACAAGTGCTATACCACTATGGTCCCACTTAGGTATCATGGTGAGACCAAAATGGTGCAAGCAGCCTTGACCCCCGATTCTTGCTAGTAA

### >His-CH2-CH3-Tp

MDAMKRGLCCVLLLCGAVFVSPAGAHHHHHHGSCQPSLSLQRPALEDLLLGSDASITCTLNGLRNPEGAVFTWEPSTGKDAVQKKAVQNSCGCYSVSSVLPGCAERWNSGASFKCTVTHPESGTLTGTIAKVTVNTFPPQVHLLPPPSEELALNELLSLTCLVRAFNPKEVLVRWLHGNEELSPESYLVFEPLKEPGEGATTYLVTSVLRVSAETWKQGDQYSCMVGHEALPMNFTQKTIDRLSGKPTNVSVSVIMSEGDGICY*

### >His-CH2-CH3-Tp

ATGGACGCGATGAAAAGGGGGCTGTGCTGCGTGCTCCTGCTTTGCGGAGCAGTCTTTGTGAGTCCTGCGG GGGCTCACCATCACCATCACCATGGCTCTTGCCAGCCCTCCTTGAGCCTCCAACGGCCGGCGTTGGAAGA CCTCCTGCTGGGTAGCGACGCTAGCATTACGTGCACCCTGAACGGCCTGCGCAATCCGGAGGGAGCTGTG TTTACTTGGGAACCTAGTACGGGGAAGGATGCGGTGCAGAAAAAAGCGGTGCAGAACAGTTGCGGATGTT ACAGTGTCTCCAGTGTATTGCCAGGCTGTGCCGAGAGGTGGAACAGTGGCGCAAGCTTTAAGTGCACCGT TACTCATCCCGAAAGCGGAACACTTACAGGCACTATAGCGAAAGTAACTGTAAACACGTTCCCTCCCCAA GTCCATCTGCTTCCTCCCCCAAGCGAAGAACTTGCACTGAATGAACTCTTGTCTCTCACCTGCCTTGTTC GAGCTTTTAACCCAAAGGAGGTACTCGTTAGATGGCTGCATGGGAACGAAGAACTTAGTCCTGAGAGCTA TCTTGTGTTTGAGCCACTTAAAGAACCTGGCGAGGGAGCGACTACCTATCTCGTTACAAGCGTGCTTCGC GTCAGCGCTGAAACATGGAAGCAGGGTGACCAATATTCATGCATGGTGGGACATGAAGCTCTGCCAATGA ACTTTACTCAAAAGACTATAGACCGGCTTTCTGGCAAGCCTACCAATGTCTCTGTATCCGTAATAATGTC TGAGGGCGATGGTATTTGTTATTGATAG

